# mzLearn, a data-driven LC/MS signal detection algorithm, enables pre-trained generative models for enhanced patient stratification

**DOI:** 10.1101/2025.01.26.634927

**Authors:** Leila Pirhaji, Jonah Eaton, Adarsh K. Jeewajee, Min Zhang, Matthew Morris, Maria Karasarides

**Author notes:** Correspondence should be addressed to L.P.

## Abstract

Metabolite alterations are linked to diseases, yet large-scale untargeted metabolomics remains constrained by challenges in signal detection and integration of diverse datasets for developing pre-trained generative models. Here, we introduce mzLearn, a data-driven MS^1^ signal-detection and alignment method that runs from mzML files without user-set parameters. Across 15 public datasets, mzLearn detects 11,442 signals on average vs 7,100 (XCMS) and 4,655 (ASARI), with higher TP (89.0% vs 77.4% vs 49.6%) and lower FP (12.5% vs 17.3% vs 18.8%), while correcting instrument drifts across large cohorts without experimental QC samples. mzLearn detected 2,736 robust metabolite signals from 22 public studies (20,548 blood samples), enabling the development of pre-trained variational autoencoder for untargeted metabolomics. Learned metabolite representations reflected demographic data and when fine-tuned on unseen renal cell carcinoma data, improved risk stratification and overall survival predictions, while feature-importance analysis (SHAP) highlighted biologically plausible lipid and carnitine signals. By producing a consistent, high-quality MS^1^ feature matrix at scale, mzLearn paves the way for developing pre-trained foundation models for untargeted metabolomics.

---

Metabolomics, the study of low-molecular-weight metabolites in biological systems, presents a snapshot of the biochemical pathways used by different cells and tissues, with alterations often linked to disease^1,2^. In recent years, the UK biobank’s metabolomic studies of over 100,000 individuals have linked metabolite profiles with disease outcomes and aging^3,4^. GLP-1 trials further highlighted the potential of metabolic intervention in reducing cardiovascular, diabetes, and other chronic diseases^5,6^. While the Human Metabolome Database (HMDB)^7^ annotates over 220,000 metabolites cataloged in the human body, current identification procedures can reliably recognize only a few hundred metabolites, a fraction of what exists^8^. Therefore, metabolomic datasets remain largely untapped due to technical challenges of population-level detection and annotation of global or untargeted metabolomic signals^9^.

Untargeted metabolomic profiling using liquid chromatography coupled with mass spectrometry (LC/MS) can provide high-throughput measurements of thousands of metabolite signals^10^. Existing signal detection methods lack reproducibility and have high false positive and low true positive rates^11,12^. The final set of metabolomic features is highly sensitive to initial parameter choices^13^, with signal overlap between methods as low as 10%^14^. While methods have been developed to remove noisy signals from detected peaks^15^, none address the critical issue of low true positive or missing biologically relevant signals. LC/MS signal detection is further limited in large-scale studies due to instrument drifts^16^, causing molecule-specific intensity drifts^17^ and non-linear fluctuation in LC retention time (rt)^18^. These drifts result in misaligned or cross-aligned signals and erroneous downstream inferences^19^. Experimental methods have been developed to estimate instrument drift, including running quality control samples (QC) at regular intervals during extended runs^20,21^. There are no standardized protocols, however, and public metabolomics repositories often lack QC data and associated information. Moreover, even with the presence of QC samples, there is no computational method to estimate both rt and intensity drift simultaneously, and existing methods addressing them separately require significant user inputs^16,22^. As a result, while publicly available untargeted metabolomics datasets are growing rapidly, their utility for combinatory analysis remains limited.

In fields such as single-cell sequencing, standardized data processing methods have already enabled the integration of public datasets to develop pre-trained foundation models^23–25^. These models have proven invaluable by capturing generalizable patterns that support downstream prediction tasks when finetuned in smaller datasets^26^. Extending pre-trained models to untargeted metabolomics could similarly enhance metabolomic data utility and uncover clinically meaningful signals. While untargeted metabolomics data contain artifacts and chemical contaminants^27^, they also harbor a wealth of unannotated metabolites with significant biological relevance^28–30^. Combinatory analysis of large-scale metabolomics studies measured in different laboratories is essential to identifying robust metabolite features across cohorts despite noise and technical variability^11^. In addition to technical variabilities, demographic and environmental factors influence metabolomic profiles^31^. Pre-trained models built on diverse biological cohorts can further learn population-level diversity and aid in identifying disease-relevant signals^32^.

In this study, we introduce mzLearn, a data-driven LC/MS signal detection algorithm that produces high-quality, robust metabolomic signals at scale while simultaneously estimating and correcting rt and intensity drift. mzLearn requires no user-input parameters and autonomously learns the experimental characteristics of the data through an iterative, data-driven process and estimates and corrects instrument drifts even in the absence of experimental QC samples. Compared to existing methods, mzLearn detects a significantly larger number of LC/MS signals with superior quality. mzLearn’s streamlined process enabled the analysis of 20,548 blood-based untargeted metabolomics datasets from diverse studies, facilitating the development of pretrained generative models for untargeted metabolomics. These models produced metabolite representations that captured demographic variations and improved downstream prediction tasks when tested on the baseline serum metabolomics data of clear-cell renal cell carcinoma patients (ccRCC). Finally, we show that the learned metabolite representation facilitated joint and adversarial learning to perform complex tasks, including prognostic and predictive stratification. Our models match and potentially surpass current gold standard, clinical grade risk criteria for predicting overall survival and can stratify patients for immunotherapy response based on baseline metabolic profiles. Available at http://mzlearn.com/, mzLearn democratizes using untargeted metabolomics datasets that can pave the way for developing foundation metabolomics models.

## Results

### method overview

mzLearn is a scalable, non-parametric algorithm for detecting, aligning and normalizing ion-level MS^1^ signals measured in untargeted LC/MS measurements. It dynamically adapts to diverse datasets by learning parameters directly from the data through an iterative, data-driven process. This enables seamless scaling to thousands of samples while addressing retention time (rt) and intensity drifts caused by extended run orders and batch effects without requiring experimental QC samples (**Fig. 1a**). We evaluated mzLearn’s performance and scalability by analyzing a total of 36 publicly available datasets with various instrumental settings (**Supplementary Table 1 and Supplementary Table 2**). Of these, 15 datasets containing targeted metabolomics features served as benchmarks for comparing mzLearn’s signal detection performance against existing methods. mzLearn consistently demonstrated superior signal quality by capturing true positives while minimizing noise. Six of these 36 datasets, each comprising over 150 samples with experimental QC samples available, termed normalization evaluation datasets, were used to validate mzLearn’s ability to estimate and correct instrument drifts without requiring experimental QC inputs. Finally, 22 of the 36 datasets, referred to as pretraining datasets, included serum untargeted metabolomics measured under the same chromatography setting. The pretraining datasets demonstrated mzLearn’s capability to integrate diverse datasets, detect metabolite signals that remain robust across biospecimens despite technical variability, and support the development of pre-trained generative models for untargeted metabolomics (**Fig. 1b**). The pre-trained learned metabolite representations effectively capture population-level variations driven by demographic factors such as age. The metabolite representations are transferrable and enhance fine-tuning performance, facilitating downstream biological discoveries, including survival analysis and clinical outcome predictions. (**Fig. 1c**).

**Table 1:** Fine-tuned VAE models with transfer learning outperform both traditional models and VAE models with random initialization. The VAE models were evaluated across binary classification, multi-class classification, and survival analysis tasks. Performance metrics included AUC for binary classification, F1 score for multi-class classification, and C-index for survival analysis. Traditional machine learning (ML) models, such as logistic regression for classification and L1-penalized Cox regression for survival analysis, served as baselines for comparison. Fine-tuned VAE models with transfer learning consistently outperformed both traditional ML models and randomly initialized VAEs.

| Performance on unseen test set | Number of test samples | Task type | Metrics | Fine-tune VAE model with transfer learn | VAE model with random initiation | Traditional ML models |
| --- | --- | --- | --- | --- | --- | --- |
| **Poor vs favorable IMDC groups** | 57 | Binary classifier | AUC | **93.48** | 82.86 | 91.78 |
| **Poor, favorable, and intermediate IMDC risk groups** | 143 | Multi-class classifier | F1 Score | **59.55** | 54.78 | 58.29 |
| **Overall survival** | 149 | Survival tasks | C-index | **67.39** | 64.45 | 64.61 |

**Figure 1:**
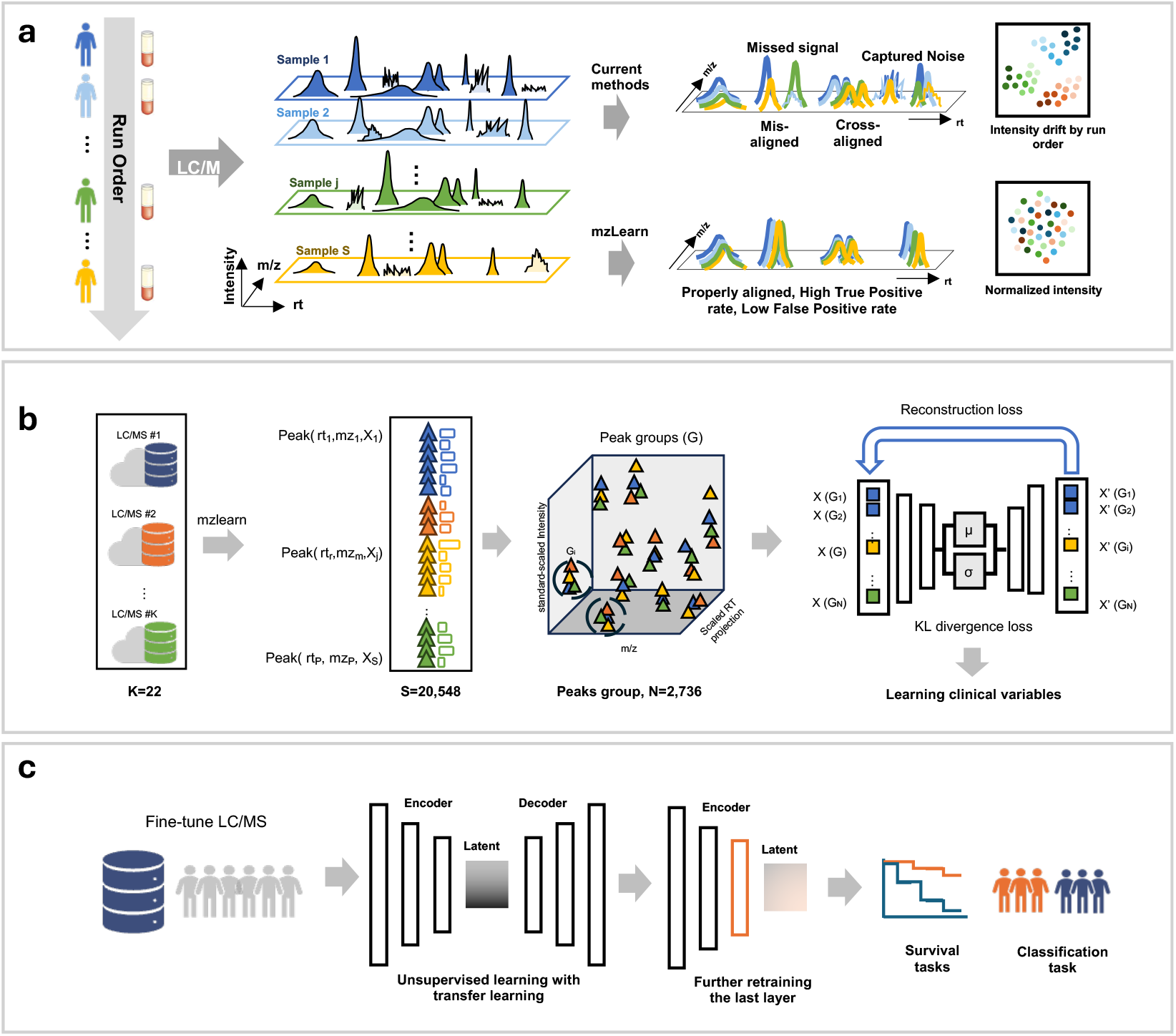
Method overview. **a**, Large-scale LC/MS measurements lead to retention time (rt) and intensity drifts due to extended run time. The current LC/MS signal detection method fails to estimate these drifts accurately, leading to missed aligning or cross-aligning signals. They further suffer from low true positives by failing to detect true signals and high false positives by capturing noise. **mzLearn** overcomes these challenges without requiring user-defined parameters. It produces high-quality signals that are accurately aligned across large datasets, with intensities normalized to account for run-order drift. This ensures the generation of robust, normalized data suitable for downstream analyses. **b**, mzLearn enables the combinations of 22 publicly availed datasets, identifies 2,736 peak groups robustly measured across studies and the development of a pretrained VAE model by minimizing reconstruction and KL divergence loss. **c**, The pre-trained models can be utilized through transfer learning to develop fine-tuned VAE models for previously unseen datasets in an unsupervised manner. These fine-tuned VAEs can be further retrained to enhance the prediction of survival or classification tasks.

### mzLearn data-driven LC-MS signal detection

mzLearn processes raw LC/MS data, consisting of data points defined by their retention time (*rt*), mass/charge (*m/z*), and intensity values^33^, stored in the open-source mzML format^34^. An individual peak is a subset of these points with specific *m/z*, rt, and intensity values. In contrast, a peak group (or feature) is a collection of individual peaks corresponding to the same underlying ion across multiple samples^35^. mzLearn’s output is a two-dimensional table of feature values defined by median rt and m/z values with normalized intensity across samples (**Fig. 2a**). mzLearn recursively partition the data into patches, which overcomes the memory constraints of signal extraction across large data sets (**Fig. 2b**). The patches are then dynamically resized by minimizing the gaps between adjacent points in rt and m/z space, which allows mzLearn to adapt to the shape of the underlying signal, reducing the need for pre-selected input parameters or prior assumptions about the peak shapes (**Fig. 2c)**.

**Figure 2:**
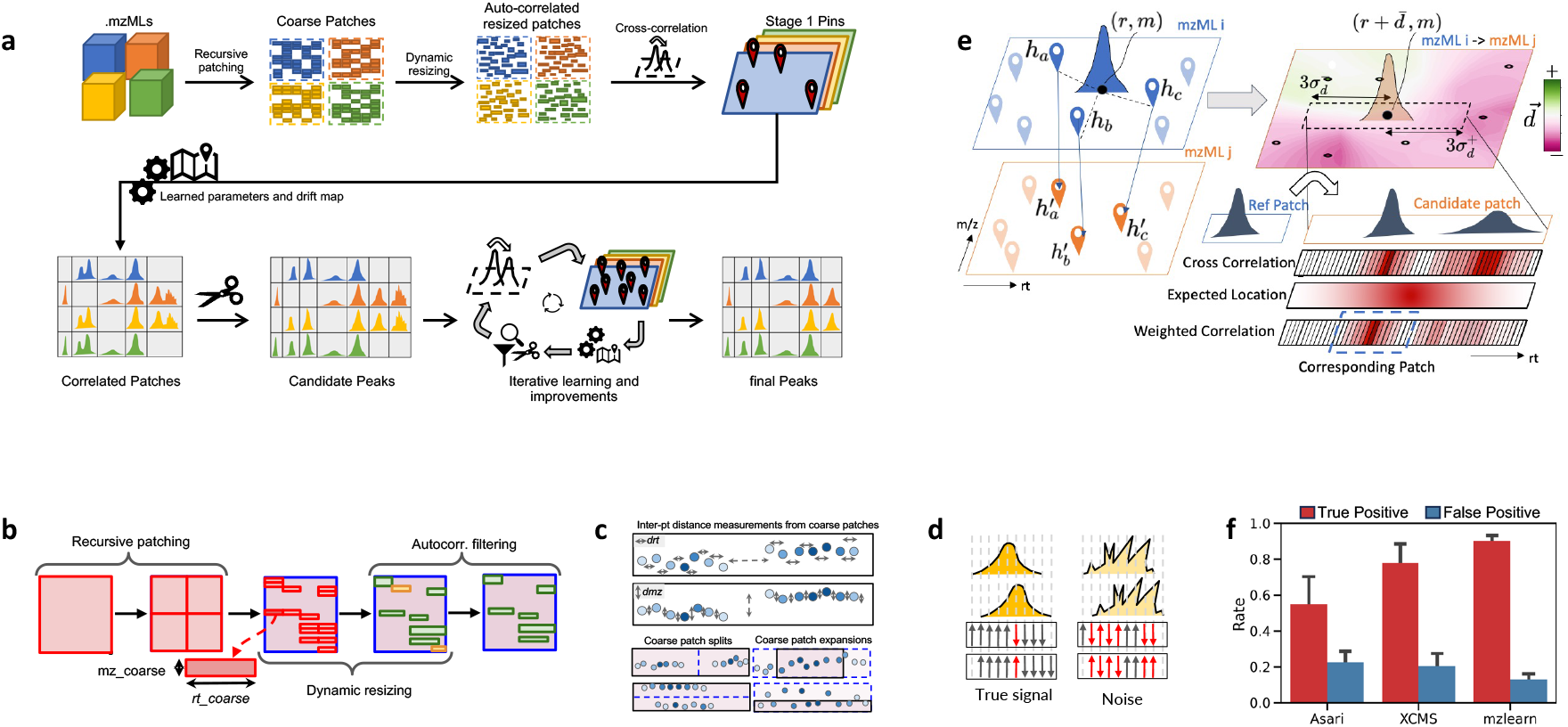
mzLearn algorithm. **a**,.mzML files are recursively partitioned into data patches, which are dynamically resized and filtered to remove noise using autocorrelation. These patches are then cross-correlated across different files to ensure alignment. Stage 1 pins, a subset of data patches with the highest auto- and cross-correlation scores, are identified to guide parameter learning. Candidate peaks are extracted from these correlated patches through a self-improving iterative process. With each iteration, the number of pins increases, improving parameter estimation and ultimately resulting in a high-quality final peak list. **b**. Each.mzML file is recursively subdivided into coarse data patches, which are dynamically resized, split, and combined to form bounding boxes around the underlying signals. These patches are then filtered using autocorrelation to remove noise. **c**, Patches are dynamically resized by analyzing the distances between individual data points. Large gaps are used to split patches, creating smaller, distinct regions, while data points with sufficiently small separations are combined into larger patches. **d**, Auto-correlation of a patch including true signal (left) and a patch including noise (right) with time-shifted versions of themselves; arrows represent an increase (up) or decrease (down) of intensity as time-shift increases (left to right). **e**, mzLearn corrects for rt drift by using high-quality signals or pins to estimate the drift between pairs of samples at any coordinate in rt and m/z space (green-red map). These estimates guide the cross-correlation between the reference patch from sample *i* (blue) to the candidate patch in sample *j* (orange). Optimal alignment is determined by the convolution of the local cross-correlation information with the global pin-based drift map information. **f**, mzLearn outperforms ASARI and XCMS in average true positive (TP) and false positive (FP) peak detection across fifteen public datasets acquired using diverse LC and MS instruments. The bar plot illustrates the average TP (red) and FP (blue) rates for each algorithm, with standard deviations as error bars.

mzLearn distinguishes true signals from noise by 1) auto-correlating the signal with a time-shifted version of itself and 2) cross-correlating the signal with different files, combined with an iterative parameter-learning approach. In the auto-correlation step, mzLearn identifies patterns indicative of true peaks by calculating the similarity of each signal patch to its time-shifted counterpart. Signal patches with high autocorrelation are retained, while those with low autocorrelation are filtered out (**Fig. 2d**). The cross-correlation procedure aligns the signal across different files by sliding a reference patch over a larger candidate region until the optimal overlap between the two regions can be found (**Fig. 2e**). Correlated patches may contain several candidate peaks. These candidate peaks are extracted from correlated patches through a multi-step scissoring process by first splitting them based on the gaps in *rt*-*m/z* space and then at the valleys between intensity local maxima (**Supplementary Fig. 1**, Online Methods). mzLearn learns the parameters of the above steps through an iterative process. In each iteration, mzLearn defines a subset of the highest-quality correlated patches, termed “pins,” which are used to estimate the auto- and cross-correlation thresholds (**Fig. 2a**). Initially, more conservative thresholds are used due to limited information. As the quality of detected peaks improves with each iteration, the number of pins increases, enabling more accurate estimation of thresholds and drift corrections.

mzLearn overcomes rt drifts in large-scale metabolomics studies by calculating an rt-drift map. The drift in retention time between two files is quantified by calculating the weighted average and standard deviation in the rt-drift among neighboring pins (**Fig. 2e**). The rt-drift map defines the bounding box of cross correlation between file pairs, where the data patches with the highest cross-correlation within the defined bounds are aligned (**Fig. 2e**). By updating thresholds and drift maps in each iteration, mzLearn searches for potential missing signals that may have been overlooked in previous steps (**Supplementary Fig. 2**, Online Methods). mzLearn converges after three iterations, producing a final high-quality list of features with minimal missing values and features aligned across thousands of samples.

We evaluated the performance of mzLearn in detecting true positive and false positive signals against XCMS^11,36^, the most widely used method among parameter-intensive approaches such as mzMine^37^ and MSDial^38^, and Asari^39^, a newer approach that minimizes reliance on key parameters. Using the benchmarking datasets, we evaluated these methods in terms of the number of identified peaks, at least in the 20% of input samples, their true positive (TP) rate, defined as the number of targeted metabolites present in the samples found by each method, and their false positive (FP) rates, where we manually labeled 100 randomly selected peaks as “good” or “bad” signals based on peak shape, signal to noise ratio, and peak alignment (**Supplementary Fig. 3**). We demonstrated that mzLearn outperformed other methods in both signal quantity and quality. On average, mzLearn identified the most LC/MS signals (N=11,442) compared to XCMS (N=7,100) and ASARI (N=4,655), absolute feature counts per dataset and associated database details are listed in **Supplementary Table 3**. Importantly, mzLearn achieved the highest signal quality, with an average true positive (TP) rate of 89.0% and a false positive (FP) rate of 12.5%, outperforming XCMS (77.4% TP, 17.3% FP) and ASARI (49.6% TP, 18.8% FP) (**Fig. 2f, Supplementary Table 3**). Per-study feature–intensity matrices with feature-level metadata (detection frequency, median m/z, median RT) are provided as Supplementary Data.

### mzLearn corrects signal intensity drift in large-scale studies

mzLearn generates synthetic QC samples and calculates feature-specific intensity drift maps to mitigate intensity drift often caused by extended run orders and batch effects. Synthetic QCs are generated through an iterative process, where samples are clustered based on both their low-dimensional intensity representations and run order, allowing only consecutive samples to form clusters. In each iteration, the synthetic QC is defined as the average intensity of each cluster (**Fig. 3a**), and an intensity drift map is created by interpolating the drift of neighboring pins in rt and m/z dimensions to estimate missing values (**Fig. 3b**). Given the diverse chemical properties of metabolites, each feature exhibits a unique drift pattern; therefore, the peak intensities are normalized against the same peak identified in the nearest synthetic QC in the run sequence. The iterative process stops when additional clusters can no longer be consistently formed within consecutive sample orders (**Fig. 3a**).

**Figure 3:**
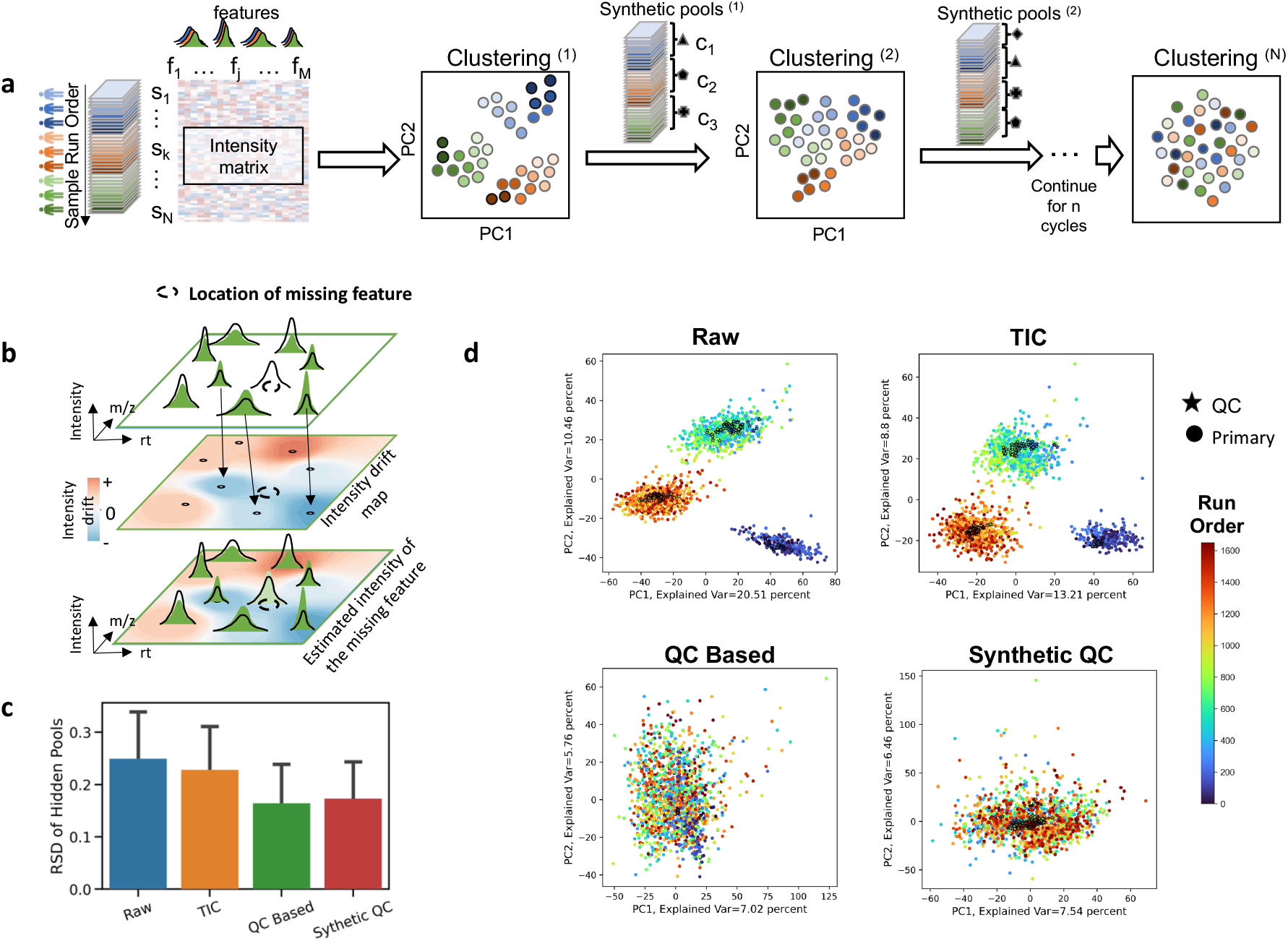
mzLearn ameliorates intensity drift. A Synthetic QC normalization overcomes signal drift. The synthetic QC samples are identified as iterative processes and normalization in each iteration. The operation is stopped where no run order-based effect on sample clustering is observable. **B**. For missing signals in QC or synthetic QC samples, the feature-specific intensity drift is estimated by interpolating the intensity drifts of neighboring pins. **d**, 4013 peaks identified in ≥60% of samples in the combined dataset of ST001236 and ST001237 were used in the following PCA plots. Each dot represents a sample, colored according to sample run order. Stars denote experimental QC samples present in the original data. PCA of the raw data without any normalization shows pronounced batch and run order effects. TIC normalization does not meaningfully address the batch effects in the data. Both the QC- and synthetic QC-based normalization applied by mzLearn largely eliminate batch and run-order effects, where QC and synthetic QC samples are now clustered closely together, and individual samples are intermixed based on their run order.

We assessed the synthetic QC normalization method in overcoming signal intensity drifts using the normalization evaluation datasets, where we withheld a subset of experimental QC samples and calculated the relative standard deviation (RSD) of peak intensities in the hidden QC sample. Synthetic QC normalization performance (17% RSD) is comparable to the performance where the peak intensities are normalized using experimental QC samples (16% RSD). It further greatly outperforms total ion concentration (TIC) normalization (23% RSD), a standard data-driven normalization method with the absence of QC samples^40^, as well as raw unnormalized data (25% RSD) (**Fig. 3c, Supplementary Table 4**). Specifically, we showed that mzLearn can identify peaks from a large cohort of ccRCC patients in Phase I (n=91) and Phase III (n=741) clinical trials (Metabolomic Workbench ids: ST001236 and ST001237)^41,42^, where the serum metabolomics data were measured in 3 distinct batches over 3 months. mzLearn successfully processes samples from two trials, combined 1,650 serum samples, detecting 4,016 untargeted features in ≥60% of samples, estimates, and corrects rt (**Supplementary Fig. 4**) and intensity drifts (**Fig. 3d**).

### mzLearn enables the combination of public untargeted metabolomics datasets to develop pre-trained generative models

mzLearn enables the development of pre-trained generative models for untargeted metabolomics by integrating data from the pretraining dataset of 22 public LC/MS serum untargeted metabolomics conducted with hydrophilic interaction liquid chromatography or HILIC in positive ion mode in several laboratories (**Supplementary Table 5**). Initially, LC/MS signals from each study were extracted by mzLearn and normalized individually to address intensity drift and intra-study batch effects. To further reduce batch effects across studies, the datasets were normalized using standard scaling. (**Fig 4a**). Next, we aligned peaks across studies using m/z values, scaled rt projections, and standard-scaled intensities (**Fig. 1b**). While m/z values are comparable between studies, rt values could be dramatically different because of the various experimental settings. The scaled rt projection between studies was learned using the combination of subalignment and graph aggregation, Eclipse^43^, and non-linear spline fitting between rt values, metabCombiner^10^ methods (**Supplementary Fig 5** and online method). We calculated a robustness score to assess measurement consistency, filtering out low-scoring peaks, and identified 2,736 robust peak groups across 22 datasets (**Supplementary Fig. 6**). Only 155 targeted metabolites (5.6% of untargeted features) were consistently detected, spanning diverse classes like amino acids, lipids, nucleotides, energy metabolites.

**Figure 4:**
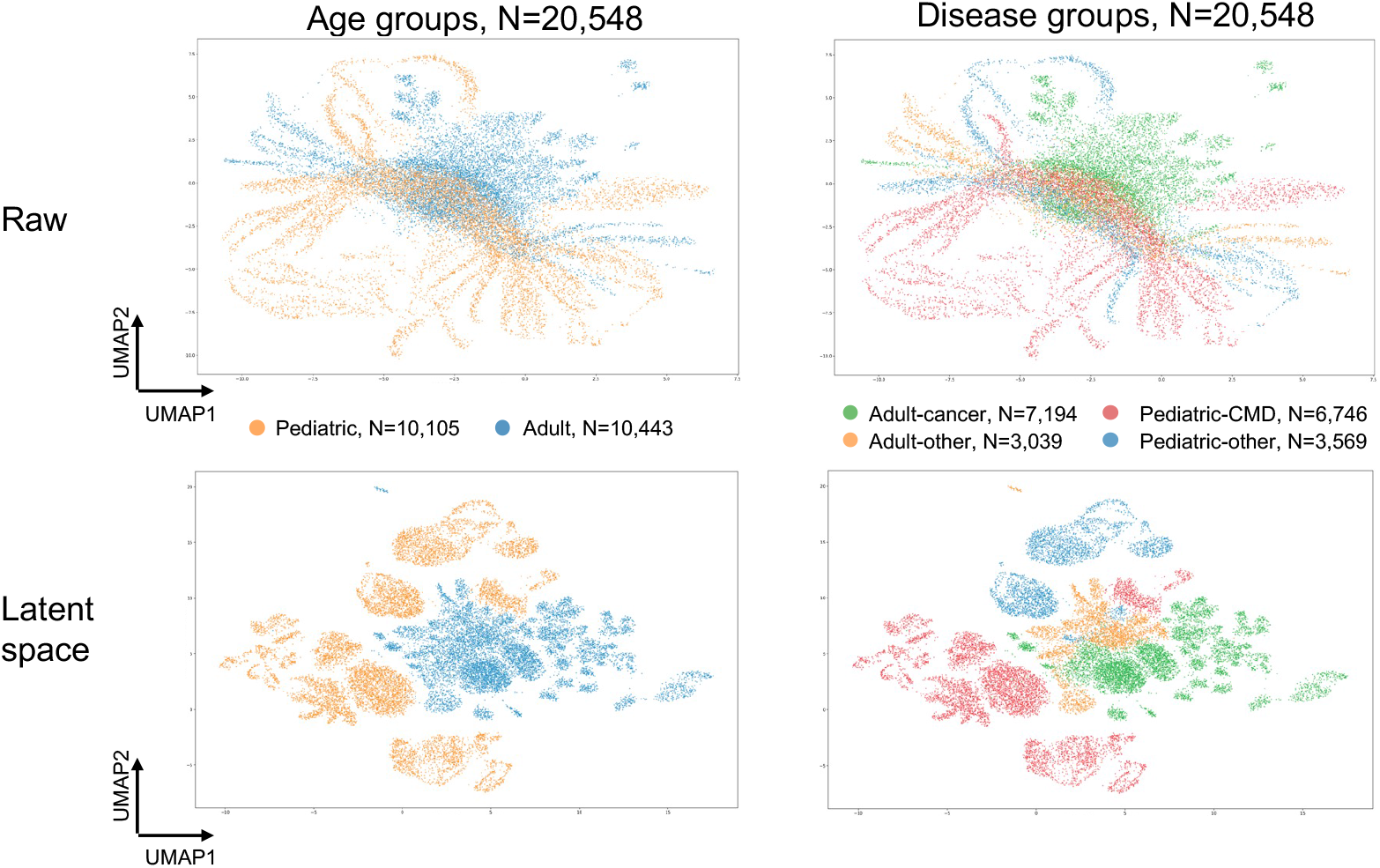
Latent space visualization of pre-trained VAE models. UMAP visualization of the raw input data (n=20,548 samples), color-coded by age and disease groups, where the samples appear mixed across these categories. Visualization of the latent space from the pre-trained VAE models, where samples naturally separate by age and disease groups, indicating that the model has captured meaningful biological distinctions.

We then learned metabolite representation by developing an unsupervised, pre-trained Variational Autoencoder (VAE) model that optimizes reconstruction and KL divergence losses^44^.VAE is a generative modeling approach^45,46^ that learns compact, latent representations of identified peak groups or metabolite signals across trained on the mzLearn MS^1^ feature matrix obtained from 20,548 blood-based samples using mini-batch stochastic gradient descent (**Fig. 1b**). Although exact metadata for each public study were unavailable, we manually labeled the studies based on disease categories and age groups of the samples (**Supplementary Fig 7**). The UMAP plot of raw data shows the mixing of samples based on age and disease labels (**Fig. 4**). Interestingly, even without supervision, the VAE model learned representations clustered samples with similar disease and age characteristics in the latent space (**Fig. 4**). While the standard-scaled intensities of the input data exhibited mixing of samples based on study labels, in the latent space, samples became further separated according to their disease and age labels (**Supplemental Fig. 8**). We further attempted to mitigate batch effects during pre-training using adversarial learning. However, due to the diverse disease profiles across studies, this approach reduced fine-tuning performance by compromising the representation of biologically meaningful features (data not shown). Depending on the specific goals of fine-tuning, the pre-trained models can be further reparametrized using a subset of samples with available metadata to create models tailored to specific clinical variables, such as gender **(Supplemental Fig. 9**, online method). This adaptability supports the model’s use across diverse clinical applications.

### Transferable metabolite representations enable fine-tuned VAE models to outperform traditional and randomly initialized models

We assessed the transferability of pre-trained metabolite representations on downstream performance by fine-tuning a VAE model using unseen baseline blood metabolomics data of ccRCC patients in the CheckMate025 Phase III clinical trial^42^, where patients were treated with an ICI (n=392) and an mTOR inhibitor (n=349). We split the samples to train (n=543), validate (n=149), and test (n=149) sets, ensuring that treatment groups, as well as clinical variables such as risk groups, age, gender, portioner therapy, etc., were balanced within each set (**Supplementary Table 6**). The fine-tuned VAE was developed unsupervised using transfer learning, while a randomly initialized VAE was created for comparison. We used Optuna, which employs a Bayesian optimization framework^47^, to fine-tune hyperparameters. Notably, the fine-tuned VAE model using transfer learning reached optimal parameters within fewer trials (n=38) compared to the randomly initialized model (n=43) **(Supplementary Fig. 10)** and achieved lower validation loss with fewer training epochs **(Supplementary Fig. 11)**.

We developed these models in an unsupervised manner, enabling rapid adaptation to new clinical questions without the need for retraining the entire model. To make specific predictions, we retrain only the final layer of the encoder and the prediction head following the latent space (**Fig. 1C**). We evaluated model performance on three distinct task types: binary classification, multi-class classification, and survival analysis. For classification tasks, we predicted IMDC (International Metastatic RCC Database Consortium) risk groups^48^, where the binary classification involved distinguishing “poor” versus “favorable” risk groups, while the multi-class classification provided a more granular prognostic categorization of “poor” vs. “intermediate” vs. “favorable.” For survival analysis, we predicted overall survival (OS) across both treatment arms, assessing model performance using the concordance index (C-index). Fine-tuned models with transfer learning consistently outperformed randomly initialized and traditional machine learning models (**Table 1**). We also explored retraining additional layers; while these adjustments improved performance on the validation set, they reduced performance on the test set (data not shown), further demonstrating that pre-trained models enhance generalizability and reduce the risk of overfitting when applied to smaller fine-tuned datasets.

### Prognostic and predictive patient stratification via joint and adversarial learning

We further evaluated whether learned metabolite representation could facilitate the development of advanced architectures for prognostic and predictive patient stratification. To develop a prognostic model capable of identifying features associated with a patient’s overall survival (OS) independent of treatment, we re-trained the final layer of the fine-tuned VAE and added a task-specific layer. We implemented two models: one trained exclusively on baseline data from patients treated with immune checkpoint inhibitors (ICIs) and the other on patients treated with mTOR inhibitors. These models were trained jointly, while the training objective combined each model’s loss, optimizing the total loss to strengthen learning across both tasks **(Fig. 5a)**. This joint training setup enables the models to learn OS-associated patterns in a treatment-independent manner, resulting in a prognostic model. We further evaluated the performance our prognostic models against the IMDC groups, established clinical grade parameters that define RCC patient sub-groups into variable prognostic risk groups^48^. Based on the predicted OS from our models, we stratified samples into poor, intermediate, and favorable risk groups, matching the number of patients in each category to the IMDC-reported distributions for a fair comparison. When tested on unseen data (n=143), the prognostic VAE models significantly outperformed the clinically defined IMDC risk groups in predicting patient OS. While only the IMDC poor versus favorable groups showed a statistically significant difference in OS, all risk groups predicted by the prognostic VAE demonstrated significantly distinct OS distributions and improved hazard ratios **(Fig. 5b)**.

**Figure 5:**
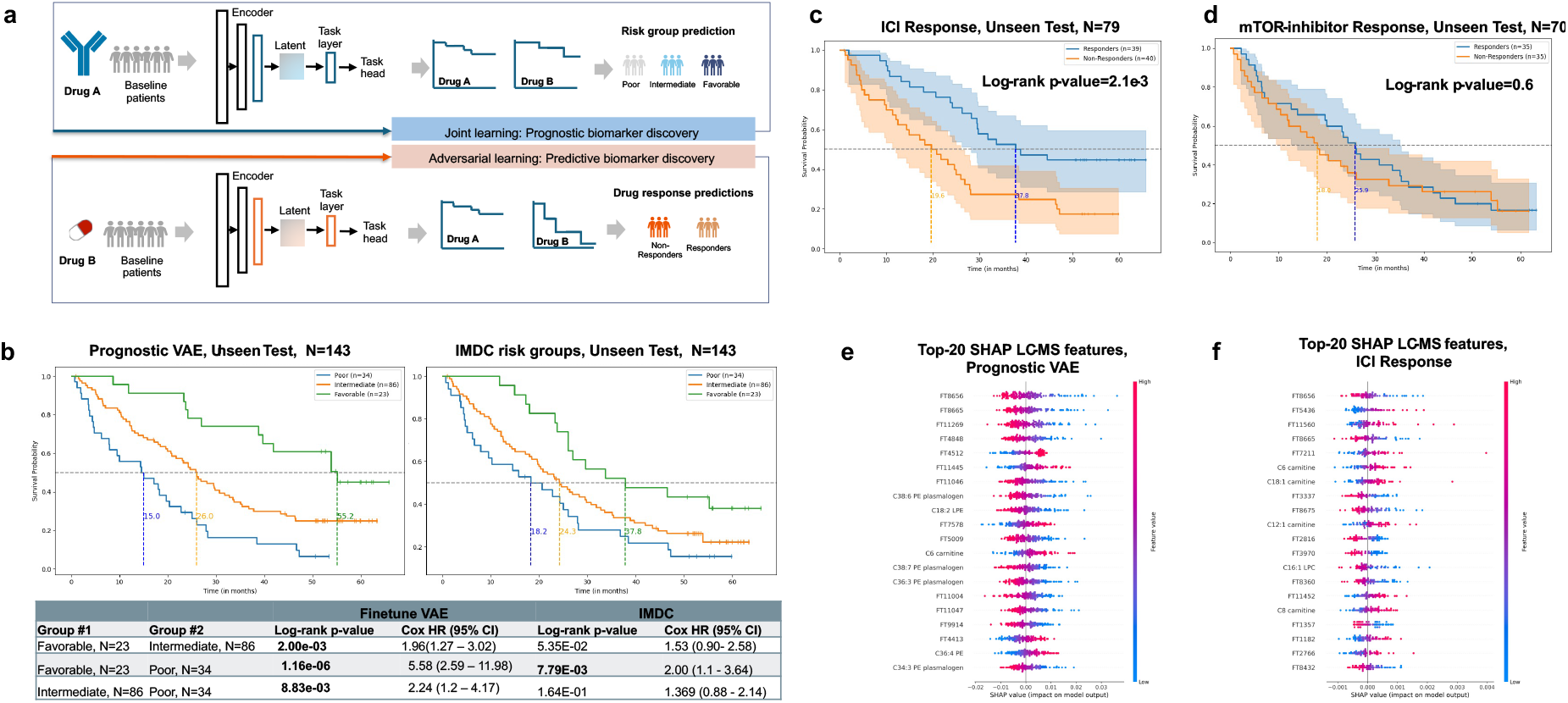
Join and adversarial learning to develop prognostic and predictive models. **a**, Joint learning approach to retrain the last layer of two fine-tuned VAE models, developed through transfer learning, with the addition of a task-specific layer to predict overall survival (OS) for Drug A (ICI) and Drug B (mTOR inhibitor). Both models jointly learn features associated with OS that are independent of treatment, forming a prognostic model. Patients are subsequently stratified into three risk groups (favorable, intermediate, and poor) based on the predicted OS. In contrast, a predictive model specific to Drug A is developed through adversarial learning. Here, one VAE model learns to predict OS for Drug A while competing with another VAE model that predicts OS for Drug B, isolating features unique to Drug A’s response and reducing confounding effects. Patients with above-average predicted OS are considered responders, while those below average are classified as non-responders. **b**, Performance evaluation of the prognostic model on an unseen test set, N=143. The prognostic VAE model outperformed clinical-grade IMDC risk groups (a standard prognostic system for ccRCC), effectively refining risk stratification based on OS predictions. **c**, Evaluation of the ICI-specific predictive model on an unseen test set, N=79, demonstrating exclusive stratification of responders to ICI with a log-rank p-value of 2.1e-3 **d**, The adversarial model cannot accurately predict OS in the mTOR arm, on an unseen test set, N=70 (log-rank p-value=0.61). e,Top-20 SHAP ion features of Prognostic VAE (treatment-agnostic OS risk across all patients). f, Top-20 SHAP ion features of ICI-specific predictive model. Beeswarm plots show SHAP values computed on held-out baseline samples, using the training set as the background. Features are ordered by mean absolute SHAP values within each model. Each dot is one patient; color encodes the measured feature value (red = higher, blue = lower), and position on the x-axis indicates the feature’s impact on the model output. Positive SHAP increases the model score (e: higher prognostic risk; f: higher probability of nivolumab response); negative SHAP decreases it. Labeled metabolites are annotated features, while the rest are untargeted LC/MS features.

We also developed an adversarial VAE model to exclusively predict overall survival (OS) for ICI therapy. As for the prognostic model, we re-trained two VAE models: one to predict OS for ICI response and the other as an adversarial model for OS in mTOR-treated patients **(Fig. 5a)**. While the main VAE aims to capture features specific to ICI response, the adversarial model prevents it from learning features associated with OS independent of the treatment. Therefore, the predictive VAE model captures drug-specific effects, reducing confounding variables and isolating biomarkers that are uniquely predictive of ICI response. We evaluated the model’s performance on unseen test data (n=143). Using the predicted OS from the predictive VAE, we labeled patients as responders if their OS was above the cohort average and non-responders if below average. The predictive VAE was trained with an adversarial objective to learn ICI-specific survival signal while suppressing any representation that also predicts outcomes in the everolimus (mTOR) arm. Accordingly, the Accordingly, Kaplan–Meier (KM) survival analysis showed significant stratification in the for ICI-treated patients (log-rank p-value = 2.1e-3, **Figure 5c**) but no stratification in for mTOR inhibition (log-rank p-value = 0.61, **Figure 5d**)

To identify the most influential metabolic features for each VAE model prediction, we computed SHAP values^49^ on held-out baseline samples using a training-set background and ranked features by mean SHAP values to capture global importance and used the cohort-average signed SHAP to assign direction relative to the model’s output (protective vs harmful). We then formed Top-20 lists per model, prognostic VAE across all patients (**Figure 5e**) and ICI-predictive VAE within Nivolumab arm (**Figure 5f**). While the majority of Top-20 features are unannotated ion features, several map to targeted analytes measured in the original CheckMate025 metabolomics dataset^42^. Only three of top-20 (15%) overlap across lists, including FT8656 and FT8665, an +1.003 Da isotopic pair at identical RT (i.e., the same underlying molecule) and FT4553 (C6 carnitine). The remaining Top-20 features are model-specific. The prognostic list contains three phosphatidylethanolamine (PE) plasmalogens (C38:6, C38:7, C36:3) and one polyunsaturated Lyso-PE (LPE, C18:2), whereas the ICI predictive model list contains four acyl-carnitines (C6, C8, C12:1, C18:1) and Lyso-phosphatidylcholine (LPC, C16:1). Together, these patterns indicate complementary baseline metabolic programs with only a small, shared core signal.

## Discussion

mzLearn is a deterministic, non-parametric algorithm that iteratively learns internal thresholds and drift maps from the data and addresses key challenges in LC/MS signal processing by offering a streamlined, user-friendly method that requires no input parameters or metabolomics expertise. mzLearn was tested on 36 public datasets with diverse experimental settings (i.e., revered-phase, HILIC, Orbitrap, QTOF, in positive and negative modes) and consistently outperformed current algorithms in the highest number of detected MS^1^ LC-MS peaks, achieving the highest true positive rate and maintaining the lowest number of false positives, while effectively mitigating instrument drifts. Existing LC/MS signal detection methods often rely on assumptions about peak shape and require numerous input parameters^50^, making them highly sensitive^39^ with non-reproducible outcomes across different algorithms^51^. While combining peak lists from multiple methods has been shown to increase the true positive rate^52^, this approach can be cumbersome and still fails to address fundamental issues of low true positives and reproducibility. In contrast, mzLearn dynamically adapts to varying peak shapes, enabling the detection of over 60% more signals (on average) compared to existing methods. Unlike other algorithms, where a higher number of detected signals has been shown to produce higher false positives^53^, mzLearn produces high-quality signals as it iteratively refines signal characteristics and filters out the noise to identify the missing true peaks.

Instrument drifts in large-scale metabolomics studies, caused by extended run orders and batch effects, significantly hinder untargeted signal detection^16^. Large-scale LC/MS experiments spanning months often lead to substantial retention time (rt) and intensity drifts, obscuring biologically relevant features. mzLearn tackles these challenges by generating synthetic QC samples and constructing intensity and rt drift maps through a dynamic, iterative, data-driven learning process that continuously estimates and corrects instrument drifts. By eliminating the need for experimental QC inputs, mzLearn streamlines workflows and enables the robust and scalable analysis of large-scale public datasets.

We deployed mzLearn on 22 studies with HILIC positive mode, encompassing 20,548 blood samples, using the resulting MS^1^ion-feature matrices to pre-trained a VAE model. Pre-trained generative models using diverse metabolomics data from various cohorts are essential for learning metabolite representation associated with demographic variables across defined patient populations. When many batches are combined, stronger correction reduces site effects but can dampen true signal, whereas lighter correction preserves biology yet leaves some batch structure. Future large-scale deployments may need to incorporate batch covariates or domain-invariant regularization if needed.

Importantly, pre-trained learned metabolite representations are transferable and improve clinical and survival prediction tasks when finetuned on baseline serum metabolomics data of ccRCC patients. In our analysis focusing on the prognostic stratification of ccRCC patients, we show the fine-tuned model can match and potentially outperform the predictive value of the gold standard IMDC criteria to predict patients’ survival. This is biologically plausible in ccRCC, as previous studies have reported genes involved in metabolic regulation associated with overall survival in ccRCC patients^54^, and the loss of the frequently mutated VHL activates hypoxia-inducible factor (HIF) signaling^55^ that rewires lipid metabolism, suppresses fatty acid oxidation (FAO) increasing lipid uptake/storage, and remodeling membranes^56^. Consistent with this, the prognostic model’s Top-20 ion features are enriched for PE plasmalogens (C38:6, C38:7, C36:3) and a polyunsaturated LPE (C18:2), a membrane redox axis in which plasmalogens buffer lipid peroxidation (vinyl-ether antioxidant), influence ferroptosis sensitivity^57^, and often mark a less aggressive state, while polyunsaturated lyso-phospholipids reflect active membrane remodeling^58^.

Metabolic adaptation has been associated with immunotherapy reports^42,59^, but large-scale untargeted metabolomic data has yet to be extensively utilized for cancer subtyping or patient stratification. In the fine-tuned adversarial model of ICI response, we can successfully classify ICI responders based on their baseline metabolic profiles, showing potential for further clinical application in treatment decision-making with additional validation. The Top-20 feature list is enriched for acyl-carnitines (C6, C8, C12:1, C18:1) and LPC (C16:1), indicating mitochondrial and FAO readiness plus an immunomodulatory lysolipid axis^60^. Acyl-carnitines report carnitine-shuttle flux and FAO capacity that support T-cell fitness^61^ and anti-tumor immunity under PD-1 blockade^62^, while LPCs can modulate immune tone^63^ and support CD8^+^ T-cell memory via major facilitator superfamily domain-containing 2A (MFSD2A)-mediated uptake^64^. The two Top-20 lists show limited overlap (3/20), but include C6 carnitine, suggesting a shared favorable metabolic tone that links lower baseline risk with greater ICI readiness. All results are at the MS^1^ ion level; they provide prioritized, biologically coherent candidates for downstream de-isotoping, annotation, and MS/MS confirmation.

While mzLearn offers substantial advantages, it’s important to discuss its limitations and potential future developments. mzLearn results are at the ion MS^1^ level, and it does not perform de-isotoping or MS/MS identification. Mapping ion features to MS/MS spectra and performing compound identification should be carried out with downstream workflows; integrating MS/MS into pretraining is an important direction for future work. Furthermore, while mzLearn’s memory efficiency allows it to scale to large sample sizes, tested processing 2,075 files in a single run, the iterative learning approach increases computational time, with analysis taking up to 2.5 days in a single machine. This highlights the rising computational demands and parallel processing on multiple compute nodes as datasets expand. Additionally, the current pre-trained model was developed specifically for HILIC positive mode. Because LC–MS experiments constitute distinct data modalities (matrix, LC chemistry, polarity, etc.), indiscriminate mixing can dilute signal.

While the method could be extended to other metabolomics measurements, integrating various LC types and modes requires the development of multi-modal learning. Future work includes processing larger datasets, leveraging transformer architectures, and testing models on multiple fine-tuning datasets to ensure generalizability.

Foundation models are advancing rapidly, with demonstrated success across diverse fields^65–67^, including single-cell sequencing. Like the genomics field, public metabolomics repositories are expanding rapidly; for instance, the MetaboLights database^68^ now includes 11,000 studies with over 1.7 million samples. Despite this growth, fully leveraging these data is challenging due to the absence of standardized QC samples to estimate instrument drift, the non-reproducibility of current peak-picking algorithms, and the need for manual parameter tuning across multiple methods to select peaks and correct drifts. mzLearn addresses these major technical problems that have not been previously solved in metabolomics. By enabling reliable and reproducible analysis of large-scale untargeted datasets, mzLearn facilitates the creation of generative models that support population-level research into clinically meaningful metabolites, which can be used for biomarker-driven drug discovery. Throughout this work, targeted metabolites constituted only 5% of the metabolomics features extracted by mzLearn. Increased depth of features detected by mzLearn, combined with available pathway enrichment analyses^69^ and network-based inference^70^ tools for untargeted metabolic data, could enable the discovery of novel metabolomic features associated with disease and treatment response, along with their underlying mechanisms. These advancements will further lead to foundation model development for untargeted metabolomics, positioning mzLearn at the forefront of tools for harnessing large-scale untargeted metabolomic datasets and enhancing its clinical.

## Acknowledgment

We thank ReviveMed’s scientific advisors, including professors Ernest Fraenkel, Matthew Vander Heiden, and Clary Clish, and previous employees Yen Lin and Julian Montagut.

## AUTHOR CONTRIBUTIONS

L.P., J.E., and A.K.J. designed the mzLearn algorithm. J.E. and A.K.J. implemented it. L.P. developed pre-trained and finetuned algorithms and implemented the models. M.Z. prepared the input datasets and developed the web app to run mzLearn. L.P., M.M., and M.K. interpreted biological findings and wrote the manuscript. All the authors approved the final version.

## COMPETING FINANCIAL INTERESTS

L.P. is a co-founder and shareholder of ReviveMed, Inc. M.Z. and L.P. are employed at ReviveMed. J.E., A.K.J., M.M., M.Z., and M.K. have stock options for ReviveMed Inc.

### Source Data

All the datasets used in the study are available in the public domain and detailed in Supplementary tables and Supplementary Data

**Supplementary information: Supplementary Tables 1–6, Supplementary Figures 1–11, and Supplementary Data**

### Code Availability

A beta release of mzLearn—including peak picking, alignment, and intensity-drift normalization—is available for non-profit academic use via a hosted, versioned Docker pipeline at mzlearn.com. Each submitted job authenticates, pulls a pinned image (e.g., `mzlearn/public:v4.5`), and executes with a unique `JOB_CODE`; users can set CPU count, and the input mzML directory is mounted read-only to ensure reproducibility. Pre-trained generative models and fine-tuning code are available for non-profit academic use at **GitHub (ReviveMed/mzEmbed)**.

## Methods

### The mzLearn Algorithm

The mzLearn algorithm operates on centroided LC/MS data in the open source.mzML format across N_tot_ samples in a cohort. Each sample file has a collection of data points, {*x*}, described by their retention time (*rt*), mass/charge (*m/z*), and intensity values (*u*): *x* = (*rt, m/z, u*). An individual peak is a subset of these points from a single sample file that have almost-identical *m/z* values corresponding to the same ion and whose intensities form a bell-like shape along the *rt* axis. If this ion is present in multiple samples, then its peaks across samples will typically have a similar shape^71^, nearly equal *m/z* values with small differences due to machine precision, and a non-linear drift along the rt axis. mzLearn captures an individual peak *p* from sample *k* when the data points that characterize the peak in the sample, {*x*^(*k*)^}_*p*_, are contained in the bounding box 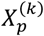. A peak group (or feature) 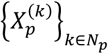 is a collection of individual peaks that correspond to the same underlying ion across *N*_*p*_ sample files. The output of mzLearn is a two-dimensional table of features, with each row pertaining to one feature that was found across *N*_*p*_ samples and each column corresponding to the intensity of the feature in those samples.

The mzLearn algorithm consists of six parts: **(A)** First, mzLearn partitions each sample file into non-overlapping dynamically resized patches that enclose all the potential peak signals in the sample and removes noisy patches via an auto-correlation process. **(B)** It then aligns the patches across samples using a cross-correlation process to form correlated-patch groups. **(C)** mzLearn estimates *rt* drift among samples and re-aligns the correlated-patch groups via the learned rt drift-map. **(D)** mzLearn extracts peaks from correlated-patch groups. **(E)** mzLearn improves and refines candidate peak groups using updated learned parameters and drift-map. **(F)** Finally, mzLearn estimates an intensity drift map and normalizes the feature intensities to correct for instrument drifts.

#### A. Creating Patches of Potential Peaks

In the first step, mzLearn partitions each sample file into non-overlapping bounding boxes that enclose all the potential peaks in the sample. We define a <u>patch</u> *X*_p_ to be one such bounding box that is dynamically resized to adapt to the shape of one underlying potential peak.

##### 1. Coarse patching

mzLearn recursively chunks the data into coarse patches that tile the retention time (*rt*) and *m/z* space for each sample file with a narrow *m/z* bin (*MZ*_*coarse*_) and a generous *rt* bin (*RT*_*coarse*_). The recursive algorithm overcomes the memory requirements for downstream analysis, reducing the computational complexity of the algorithm from O(M) to O(log(M)), where M is the number of data points {*x*} in a single LC/MS file, where *x* = (*rt, m/z, u*).

We initialize the *MZ*_*coarse*_ value to be 2 × *mz*_*tol*_ × *MZ*_*max*_, where *mz*_*tol*_ is the reported machine *m/z* tolerance in Daltons from the.mzML metadata information, and *MZ*_max_ is the maximum reported *m/z* value across all the samples. We initialize the *RT*_*coarse*_ value to be (RT_max_)/5, where RT_max_ is the maximum reported retention time across all sample files.

##### 2. Dynamic resizing of patches

Patches are then dynamically resized to ensure that each patch contains only potential peaks, and empty patches are discarded. Within every non-empty patch, points are sorted along the *rt* axis, and the distances between successive points along the *rt* and *m/z* directions are measured as *drt* and *dmz*. We define *D*_*rt*_ and *D*_*mz*_ as the acceptable lower bounds for distances between successive points in a patch along the *rt* and *m/z* axes:

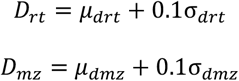

Where *μ*_*drt*_, *μ*_*dmz*_,*σ*_*drt*_ and *σ*_*dmz*_ are the mean and standard deviation of the distances between successive points along the *rt* and *m/z* axes, respectively. The inferred *D*_*rt*_ and *D*_*mz*_ allow us to adapt to different machine precisions and prevent the issue of peak fragmentation (referred to as “bleeding” in ^11^) without requiring any prior knowledge about the instruments or hard assumptions about the data. A non-empty patch is then split into two separate patches if any distance between successive points along the *rt* or *m/z* axes exceeds the *D*_*rt*_ and *D*_*mz*_ pads, while patches with either differences along the *rt* or *m/z* axes smaller than *D*_*rt*_ and *D*_*mz*_ are combined.

##### 3. Removing noisy patches by calculating Autocorrelation

Next, the noisy patches are filtered by calculating an autocorrelation score. A resized patch, *X*_p_, is correlated with a time-shifted version of itself. The *M* data points in a patch are sorted along the rt axis, *X*_p_ = {*x*_0_, *x*_5_, *x*_6_, …, *x*_7_}, such that *rt*_0_ ≤ *rt*_5_ ≤ *rt*_6_ ≤… ≤ *rt*_7_. The autocorrelation for a patch, *X*_p_, is calculated as:

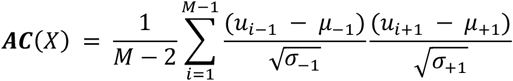

where *μ*_−1_ and *σ*_−1_ are the mean and standard deviation of the intensities for the data points *x*_0_ through *x*_+1_. Similarly, *μ*_*M*−2_ and *σ*_+1_ are the mean and standard deviation of the intensities for the data points *x*_2_ through *x*_*M*_. While a patch with a well-defined smooth peak shape will correlate strongly with itself, a noisy patch has a low autocorrelation score. We filter patches whose autocorrelation scores do not exceed a certain threshold. We seed the filter with 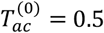 for a single pre-learning pass. After Stage-1 pin discovery and re-alignment, we set the autocorrelation cutoff to the **5th percentile** of per-peak autocorrelation among pins and **recompute it once more** after Stage-2 (three iterations total)

### B. Aligning Patches Across Samples using Cross-Correlation

mzLearn uses the cross-correlation primitive, an important concept that uses the similar peak-shape assumption to align and group peaks across samples. This operation is performed between a reference patch (X_p_) from sample k, and a candidate bounding box (X*′*) in sample k’. If the limits of the reference patch X are defined as [*rt*_*min*_, *rt*_*max*_] × [*mz*_*min*_, *mz*_*max*_], then the limits of the candidate bounding box X*′* are [*rt*_*min*_ + *lb*_*d*_, *rt*_*max*_ + *ub*_*d*_] × [*mz*_*min*_ − *mz*_*tol*_, *mz*_*max*_ + *mz*_*tol*_], where *lb*_*d*_ and *ub*_*d*_ are lower and upper bound drift estimates between samples k to k’. Initially, we use default values for the bounds: *lb*_2_ = −*RT*_*max*_/50, and *ub*_2_ = *RT*_*max*_/50. The lower and upper bound estimates are associated with the drift in rt and are updated later by the rt-drift map (Section C.2).

To calculate the cross-correlation between the reference patch (X_p_) and the candidate bounding box (X*′*), the data must be binned along the *rt* axis to form arrays of intensities. The *rt* axes in each bounding box are binned with bins of size Δ, considered as *μ*_*drt*_/10. Bins with more than one data point have their intensities averaged. If the distance between *rt*-consecutive data points is greater than *D*_*rt*_, then bins between those points are imputed with the minimum intensity value. Otherwise, the empty bins are filled with interpolated intensity using the nearby non-empty bins. The output of this step is the array *F*_Δ_(*X*)[*rt*], which is the binned interpolated data of patch X and is defined to be zero when *rt* is outside the range of *F*_Δ_(*X*).

The reference patch is aligned to the candidate bounding box at the *rt*-displacement (*d*), which maximizes the alignment function between their bounding boxes. The alignment function (**AL**) is defined as:

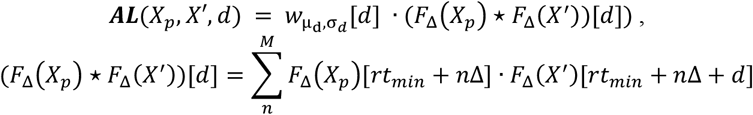

Where *d* is the *rt*-displacement between *X*_*p*_ and *X*′, *rt*_*min*_ is minimum *rt* of the reference bounding box, M is the length of *F*_Δ_ (*X*_*p*_), and 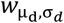 [*d*] is the expected drift function. Initially, the expected drift function 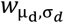 [*d*] is 1 for all values of *d*. Later, the expected drift function is determined by the *rt*-drift map (Section C.2) using the expected *rt*-drift (*μ*_d_) and the *rt*-drift uncertainty (*σ*_*d*_) between samples k to k’. With *rt*-drift map information, 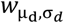 [*d*] is 1 for *d* in the 2*σ*_*d*_ neighborhood of *μ*_d_, and decays to 0 for *d* beyond this neighborhood.

The limits of the candidate patch *X*′_*p*_ are defined as [*rt*_*min*_ + *d*^*\**^, *rt*_*max*_ + *d*^*\**^] × [*mz*_*min*_ − *mz*_*tol*_, *mz*_*max*_ + *mz*_*tol*_], where *d*^*\**^ is the *rt*-displacement that maximized ***AL***(*X*_*p*_, *X*′, *d*). The candidate patch and the reference patch form a patch-group pair 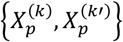. After alignment of the patches, we calculate the cross-correlation score, which is the zero-mean normalized cross-correlation, to return values between −1 and 1:

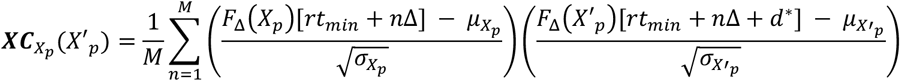

where *μ*_*X*_ and *σ*_*X*_ are the mean and standard deviation of *F*_Δ_(*X*) for patch *X*, and M is the length of *F*_Δ_(*X*_*p*_). Patch pairs above the cross-correlation threshold are kept as correlated patches. We seed with 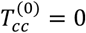. After Stage-1, the cross-correlation cutoff is set to the **5th percentile** of per-peak cross-correlation among pins and is **recomputed once** after Stage-2; these learned cutoffs replace the seeds for all subsequent filtering.

After we obtain pairs of cross-correlated patches, all pairs that share the same referential patch are combined into a cohort wide patch-group 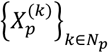, where *N*_*p*_ is the number of patch pairs.

### C. Estimating *rt* Drift

To estimate the *rt* drift between pairs of correlated patched, we define “pins” as a subset of the correlated patches found in most sample files with above-average autocorrelation and cross-correlation scores. Pins are updated at each stage of mzLearn (via updated filtering parameters and drift information) and are used for the next stage of the peak improvement process.

#### 1. Stage-1 Pins and rt-Drift Map

The stage-1 pins are patches that cross-correlate across all samples (no missing values) with initial group-average cross-correlation scores exceeding 0.8 and are used to obtain *rt*-drift map information. Stage 1 pins, the set of T robust patches h_1_, …, h_T_ spanning the entire cohort of files, are then used to generate non-linear drift maps between any desired pair of samples. Specifically, the drift 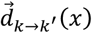 from any sample *k* to *k*^*′*^ at the desired coordinates *x* = (*rt, m/z*) can be interpolated by computing the weighted average (*μ*_d_) of the *rt*-drift of pins, where pins closer to the coordinate of interest are given a higher weight and pins further away are given lower weight. The weighted standard deviation (*σ*_*d*_) of the *rt* drift is used to measure the error tolerances in the drift. The weighted 1^st^ and 99^th^ percentiles of the pin drifts are the lower (*lb*_2_) and upper bounds (*ub*_*d*_) of the drift at *x* in sample *k*, respectively.

#### 2. Re-alignment of patches via the rt-drift map

We repeat the cross-correlation primitive for every file pair in the data set using the inferred *rt*-drift map. Unlike the cross-correlation used to find stage-1 pins, the stage-1 drift map 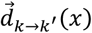 is now available to guide the cross-correlation procedure. The relevant *rt*-drift information for the cross-correlation of patch p in file k to sample k’ is obtained by considering the stage-1 drift map at the peak position of the patch:

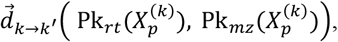

where 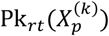 and 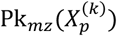 are the *rt* and *m/z* coordinates of the patch 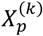 maximum intensity. The rt-drift information is then used to guide the alignment of 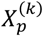 with the corresponding signal in sample k’ via the cross-correlation primitive (Section B).

### D. Peak Extraction via Multi-Step Scissors

While most patch groups include data from one peak, some patch groups may contain multiple peaks with similar *m/z* and *rt* values, for example, peaks from isomeric metabolites. To separate multiple peaks in the same patch group, we apply a procedure named “scissors” according to the following steps:

a. Patch groups are split by gaps in *rt*. The reference sample (*k*_0_) adjusted *rt* data points 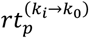 are collected from the individual patches in the patch group, 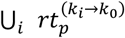. If there is a gap between consecutive *rt* values that is greater than *D*_*rt*_, the group is split at the gap.
b. Patch groups are split by gaps in *m/z*. The reference sample (*k*_0_) adjusted *rt* and *m/z* data points 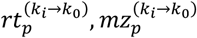 are collected from the individual patches in the patch group. If the group-wide *m/z*-width is greater than 10*mz_tol_(Da), the patch group is split by k-means clustering. The number of clusters for k-means clustering is chosen using DBSCAN, which maximizes the density of points within each cluster while minimizing the density between clusters^72^. New groups are created from each cluster of data points.
c. Patch-groups are split at the *rt* position of the local intensity minima. mzLearn searches for the *rt*-locations of the intensity minima, **Min**_***rt***_, in a patch group, 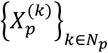, with two strategies: In the first strategy, the intensity minima are measured from the combined patch group signal, 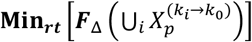. In the second strategy, the intensity minima are determined by consensus: Intensity minima are found in each individual patch, and those locations are averaged, 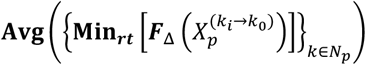.

The result of the multi-step scissor procedure is an expanded set of candidate peak-groups.

### E. Improving and Refining Peak Groups

Peak groups are further filtered to remove noisy and mis-aligned peaks, based on the improved parameters learned from Stage-2 pins, which are a subset of candidate peak groups with high quality. Here, we consider the candidate peaks that are found in 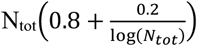 samples, where N_tot_ is the total number of samples. Features that have above-average autocorrelation scores and above-average cross-correlation scores are selected as stage-2 pins. Stage-2 pins are used to obtain a more accurate stage-2 drift map and to calculate quality thresholds.

#### 1. Alignment Reliability Score

The alignment reliability score, 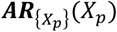 of an individual peak *X*_*p*_, with respect to its peak group, {*X*_*p*_}, as a measure of the consistency of the peak *rt*-location across all samples in the peak group. The adjusted peak coordinate, 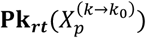, is the *rt*-coordinate of the k^th^ sample’s maximum intensity value in the reference (sample *k*_0_) coordinate system. The alignment reliability score compares this adjusted peak coordinate to the rest of the patches in the group to identify any outliers:

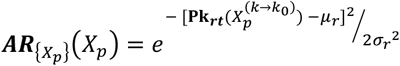

where 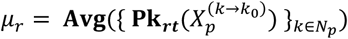 and 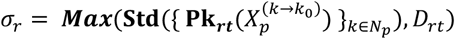. The alignment reliability score 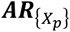 is 1 if the patch’s peak *rt*-location appears in a location consistent with the rest of the group, and 0 if the peak location is an outlier.

#### 2. Peak Quality Filtering

Stage-2 pins are used to learn the numerical thresholds for quality features. We define *g(X*) as a function that acts on an individual peak from a peak-group, 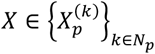, and returns a quality score between −1 and 1. We define an individual-level score threshold, *th*_V_, such that if *g(X*) < *th*_*g*_, then the peak is removed from the peak-group. We define a feature-level score threshold, *th*_{*g*}_, such that if 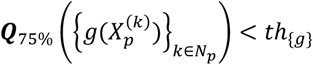, then the full peak-group is removed from the list of possible features. We learn *th*_{*g*}_ from the set of pins, H, by taking the 5^th^ percentile of the *g(X*) for all the individual peaks from the pin groups:

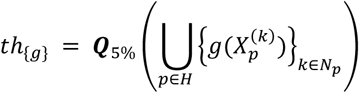

Then, the individual-level thresholds are defined from the feature-level thresholds:

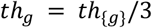

We use the above formula to define the minimum acceptable autocorrelation score for a peak-group and individual peak (*g* = ***AC***), and the minimum acceptable cross-correlation score for a peak-group and individual peak 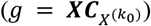, as well as the minimum acceptable alignment reliability score for an individual peak 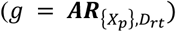.

#### 3. Missing Peak Search

After filtering, the original patches have passed through many different operations that used conservative filters to remove potential peaks and less-accurate drift maps to conduct the alignment. Therefore, the filtered peaks, while more trustworthy, have a high number of missing values due to the algorithm’s learning process. We can recover the missing peaks using the more accurate Stage-2 drift map.

We consider a peak group with individual peaks found in *N*_*p*_ samples, 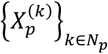, and choose a file, *k*′ ∉ *N*_*p*_, that is not included in the peak group. We survey the individual peaks from the peak group, and for each surveyed peak, we estimate the position in file *k*′ where we would expect to find the corresponding peak. The bounding box estimates from all surveyed files are averaged together to create a final estimate of the expected peak location, 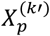, in sample *k*′. The data in expected peak location 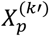 is added to the peak group if it passes the individual peak quality thresholds defined above. After the missing peaks have been added to their respective peak groups, the merging, scissor, and filtering operations discussed previously are repeated.

### F. Estimating Intensity Drift and Normalization

mzLearn is equipped with a normalization method that estimates signal intensity drift caused by extended run order and batch effects. The first step, pin normalization, removes sample-wide technical variations, such as those due to differences in the sample concentration or sample injection volume. Where no pool or QC samples are interspersed between the primary LC/MS samples in large-scale cohorts, mzLearn creates synthetic QC through an iterative process and normalizes the data in each iteration. Finally, it estimates the intensity drifts for the cases of missing values in pool or synthetic pool samples.

#### 1. Pin Normalization

Pin normalization corrects the sample-wide technical variations in the data. Peaks are normalized sample-by-sample by dividing by the total intensity of the stage-3 pins. Stage-3 pins are defined as those features found in ≥60% of the sample files with auto-correlation and cross-correlation scores above the 10^th^ percentile.

#### 2. Creating Synthetic Pool Samples

mzLearn creates Synthetic QC samples from the primary samples that approximate experimental QC samples, which can be used to estimate the feature-specific intensity drift when no QC samples are available in the public domain datasets. First, primary samples are clustered based on a low-dimensional representation of their current feature intensity profile, with each cluster constrained to include only consecutive samples in the run order. The number of clusters is chosen to maximize the Silhouette Coefficient while maintaining at least 20 samples in each cluster. Second, Synthetic pools are created by averaging the feature intensity of all samples within a cluster. Then, peaks are partially normalized to the intensity of their corresponding synthetic pool, where the silhouette coefficient of the clustering modulates the normalization factor. This clustering, synthetic QC generation, and modulated normalization process iterates until run-order consecutive clusters have a consistent near-zero silhouette coefficient.

#### 3. Feature-Specific Intensity Drift-Map

QC samples (or their synthetic equivalents) can be used to estimate the intensity drift for a specific feature. In the case when a feature, *f*, has no missing values in the QC samples, mzLearn computes the normalization of its intensity in sample file k, 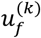, as:

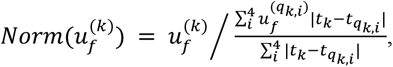

where *t*_*k*_ is the experiment run time associated with sample k, 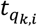 is the experiment run time associated with pool sample *q*_*k,i*_, and *q*_*k*,1_, *q*_*k*,2_, *q*_*k*,3_, *q*_*k*,4_ are the four pool samples closest to sample file k in the experiment run time. When 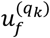 is a missing value; mzLearn estimates the value of 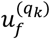 using nearby existing features in the QC sample.

mzLearn can correct feature-specific intensity drift, even when features have missing values. mzLearn considers the set of features with ≥60% sample frequency in the set of QC or synthetic QC pins, *H*_*pool*_. For each QC or synthetic QC sample, we compare the intensity of the identified features with the QC- or synthetic QC-wide mean *μ*_*f*_ to create an intensity drift map for that QC or synthetic QC sample. This intensity drift map, 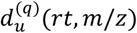, is used to estimate the intensity for the missing feature, 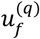 using its *rt*-*m/z* coordinates:

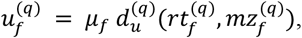

where 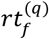 and 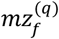 are the *rt* and *m/z* coordinates of the missing feature in the *q*^th^ QC or synthetic QC sample. The drift map 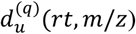 is created by interpolation using the existing (not-missing) features *h ϵH*_*pool*_ in the *q*^th^ QC or synthetic QC sample, 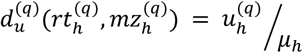.

### G. Data-driven threshold learning and update schedule

#### 1. initialization (Stage 0)

We apply a one-time seed of 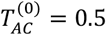, to filter obviously noisy patches and 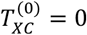, to retain non-negative cross-correlations.

##### 2. Stage-1 pins and drift map

We define pins as correlated patch groups present in all samples whose group-average cross-correlation ≥ 0.8 and whose average autocorrelation exceeds the cohort average. the 0.8 coherence anchor yields high-precision alignment landmarks; percentile bounds make the search window data-adaptive and robust to outliers. Pins generate a non-linear rt-drift map by weighted averaging of local pin drifts (weights decrease with rt–m/z distance). The lower/upper bounds of expected drift are the weighted 1st/99th percentiles and restrict the search window for cross-correlation in later stages.

#### 4. Learning thresholds from pins (Stages 2–3)

For each quality metric g∈;{AC, XC, AR}, we compute the empirical distribution of per-peak scores across the current pin set and set the feature-level cutoff to the 5th percentile. A 5th-percentile quantile policy is scale-free and controls tail noise while preserving recall; three passes are sufficient to reach a stable feature set and drift map. A candidate feature passes if its group-level score if it is more than the cutoff; a member peak is retained if its individual score is higher than the same cutoff. Thresholds are relearned after each re-alignment; mzLearn converges in three iterations.

##### 4. Normalization sets and synthetic-QC stopping

For pin normalization, we define Stage-3 pins as features found in ≥ 60% of samples whose autocorrelation and cross-correlation exceed the cohort 10th percentile. Synthetic-QC clustering is constrained to run-order-contiguous groups (≥ 20 samples), iterated until the silhouette coefficient approaches 0 (no residual run-order structure). These thresholds are conservative— high enough to avoid noisy anchors, low enough to ensure sufficient coverage for normalization and drift control.

The sensitivity analysis by varying the learned quantiles and the iteration count, the detected feature sets and alignment residuals changed minimally, and downstream conclusions were unchanged. Each run writes a machine-readable CSV (`params_overall.csv`) with instrument precision, *MZ*_coarse_, *RT*_coarse_, and global *m/z*–RT bounds/dispersion used to initialize patching and alignment; this file is deposited alongside results to document the learned and initialized settings at run time.

#### mzLearn implementation and runtime

mzLearn is implemented in Python. The detector/alignment engine uses the scientific stack (NumPy/SciPy/pandas) for correlation-based operations and ingests centroided MS^1^ from mzML via a Python parser. The software environment is provided as a version-pinned Docker image with a matching requirements.txt for reproducibility. Runtime is dominated by cross-sample correlation and iterative drift-aware re-alignment and therefore increases with the number of mzML files in a cohort; the memory footprint remains modest due to tiling and batched processing. For larger studies, we recommend multi-core parallelism and, when available, multi-node execution.

### Peak picking evaluation

To evaluate peak-picking performance across different methods, we selected a database from Metabolomics Workbench featuring diverse experimental settings and the availability of targeted metabolomic measurements where each target has recorded m/z and rt values Supplementary Table 3 provides a detailed list of datasets and their associated IDs, which were obtained from the Metabolomics Workbench and MetaboLights databases. All.raw files were centroided and converted to.mzML format using the Microsoft Windows software MSConvert. Each data set was submitted individually to mzLearn, XCMS (version 3.18) and ASARI^39^. No parameter needs to be provided to mzLearn. To run XCMS^11^, we used “Isotopologue Parameter Optimization” (XCMS-IPO)^73^ to learn the parameters of each model, and default parameters were used for ASARI.

As XCMS’s output can vary depending on the input parameters, we used “Isotopologue Parameter Optimization” (XCMS-IPO)^73^, which aims to learn the parameters using a multi-stage optimization guided by ^13^C isotope peaks and default mass resolution of 5 ppm was used for ASARI. To evaluate model performance, we focused on peaks identified by each method that met a minimum frequency threshold of 20%. Targeted metabolite peaks were identified and considered true positives (TP%) among detected peaks by comparing reported m/z and rt values from each dataset and matching them with reference values for each method.

### Benchmarking with targeted references

For datasets with targeted measurements, we used the target lists provided by the original studies (repository-reported m/z and RT) as ground truth (Supplementary Table 3). After running each method, we considered features present in ≥ 20% of samples and counted a targeted metabolite as detected (TP) when a feature matched within |Δm/z| ≤ 0.005 Da and |ΔRT| ≤ 2% of the total RT span (identical windows for all methods). False-positive (FP) rates were estimated by manual QC of 100 randomly selected detections per dataset (peak shape, SNR, alignment).

### Evaluation of the normalization

To assess the effectiveness of our normalization method in mitigating run order effects, we selected public datasets with a minimum of 150 primary and at least 20 QC samples. For each dataset, 10% of QC samples were withheld to evaluate the normalization performance. We applied three normalization approaches: Total Ion Current (TIC) normalization, QC-based normalization, and synthetic QC normalization. In TIC normalization, each sample is scaled by the total sum of the intensities of all detected ions, while in QC-based and synthetic QC normalization, each feature is adjusted based on the closest QC or synthetic QC sample in the run sequence, as described previously. After normalization, we calculated the Relative Standard Deviation (RSD) of the hidden QC samples, which serves as a metric for evaluating each normalization approach in compensating for run-order effects.

### Pretraining datasets

The pretraining data consisted of 22 publicly available datasets, including 20,827 samples, as detailed in **Supplementary Table 5**. Each dataset was manually labeled according to age groups (adult versus pediatric) and disease categories, including adult cancer, adult other, pediatric cardiometabolic, and pediatric other. Additionally, metadata was parsed for a subset of samples to capture specific information such as exact age and BMI. We included only datasets focused on blood-based metabolomics analyzed by HILIC in positive ion mode to maintain analytical consistency. Each dataset was individually processed using mzLearn, resulting in a normalized peak intensity matrix for each study.

To identify peak groups common across these cohorts, we applied retention time (RT) alignment and scaling adjustments using Eclipse^43^ and metabCombiner^10^ methods. We first selected combined ST0001236 and ST0001237 datasets as the reference study due to their high-quality peak data, characterized by high true positive rates, low false positive rates, known target information, and a larger number of detected peaks across 1,650 samples. Each pre-trained study was then aligned to the reference study using both methods. The alignment parameters for Eclipse and metabCombiner were optimized using the known targeted metabolite information.

The goal was to maximize the number of common targets retained after alignment compared to those detected before alignment. We found that a union of both methods provided the highest number of aligned targeted metabolites across studies, prioritizing alignments identified by metabCombiner when the two methods disagreed and performed best based on this metric.

We then calculated a robustness score for each peak group, factoring in the frequency of detection within each cohort, the cohort sample size, and the number of studies in which the peak was consistently detected as:

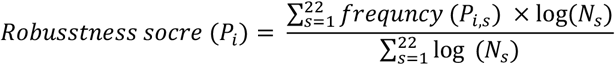

Where *P*_*i*_ is the *i-*th peak group, frequency (*P*_*i,s*_) represents the frequency of detection of *P*_*i*_ in study S, and *N*_*s*_ is the number of samples in Study S. Peaks with a minimum robustness score of 0.15 were selected, resulting in a final set of 2,736 peak groups shared across datasets. For each study, the identified peak groups were standardized using z-score scaling while missing values within were imputed by averaging.

We then split the pretraining data (n=20,548) into training (n=17,465), validation (n=2,055), and test (n=1,028) sets, where the available metadata, study cohorts, age, and disease groups were balanced across splits.

### Pre-trained VAE models

Using the pretraining data, we developed an unsupervised Variational Autoencoder (VAE) model to learn meaningful low-dimensional representations in the latent space. To ensure a balanced model architecture, hidden layer sizes were scaled according to input and latent sizes, creating an efficient structure for both encoding and decoding complex data patterns. The hidden layer size is calculated as:

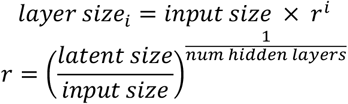

The model was trained using a composite loss function that included reconstruction loss to capture accurate data representation and Kullback-Leibler (KL) divergence loss to enforce latent space regularization:

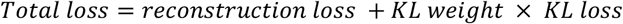

KL-annealing was applied to gradually increase the weight of the KL loss during training, allowing the model to first focus on reconstructing input features accurately before imposing latent space constraints. To prevent overfitting, early stopping was incorporated by monitoring validation loss. Additionally, the model’s hyperparameters—including learning rate, number of layers, and dropout rates—were optimized using Optuna, a Bayesian optimization framework, to enhance model performance by finding the best possible hyperparameter configuration.

To adapt the pre-trained VAE model for clinical variable representation such as age and BMI, we retrain only the last layers of the encoder, preserving the rest of the network structure and pretrained parameters. This approach allows the VAE to retain its foundational biological representations learned during pretraining while specializing in clinical tasks. In this step, we applied a targeted grid search rather than full hyperparameter optimization, focusing on a limited set of parameters, specifically the learning rate and number of layers to train in the final stage. Learning rates and layer configurations (1–2 layers) were explored to balance training efficiency with performance. This strategy allowed the VAE to effectively represent clinical attributes such as age, BMI, and disease type, providing a flexible yet reliable foundation for clinical prediction and exploration

For all model development stages, the best hyperparameters were chosen based on validation set performance. The top-performing model on the validation set was then tested on an independent test set, and the final model was selected based on its test set performance, ensuring generalizability and robustness. After training, we used the average latent space across all samples to create a UMAP (Uniform Manifold Approximation and Projection) visualization, providing an interpretable, low-dimensional data view.

### Fine-tune VAE models

We developed fine-tuned VAE models using metabolomics data from the serum of ccRCC patients at baseline (n=741, with n=392 receiving ICI and n=349 receiving mTOR inhibition). First, the baseline patients were split into training (n=443), validation (n=149), and test (n=149) sets, with clinical variables including sex, age, study region, prognosis, and prior treatment regimens balanced across splits and within treatment arms.

Then, we fine-tuned the VAE model on this dataset in an unsupervised manner, using the pretrained weights to retain the general biological representations learned during pretraining. In parallel, we developed a comparison model without transfer learning, reinitializing the VAE with random weights. In parallel, we trained a comparison model without transfer learning, reinitializing the VAE with random weights. For both configurations, we used Optuna to optimize hyperparameters, including learning rate, dropout rate, and KL weight, aiming to minimize the total of reconstruction loss and KL divergence on the validation set.

After fine-tuning the VAE on the baseline ccRCC dataset, we adapted it for three distinct tasks: binary classification, multi-class classification, and survival analysis. For each task, only the last layer of the fine-tuned VAE was retrained to retain core latent representations, with a grid search used to tune the learning rate. The model configurations achieving the highest validation AUC for binary classification, F1 score for multi-class classification, and C-index for survival analysis were selected. These top-performing models were then evaluated on an unseen test set to compare the effectiveness of VAE models with and without transfer learning against traditional machine learning models.

#### Joint Learning for Prognostic Model Development

To develop a robust prognostic model for identifying features associated with overall survival (OS) independent of treatment, we used two fine-tuned VAE models that had already undergone unsupervised fine-tuning on baseline ccRCC data. For each treatment group—immune checkpoint inhibitors (ICIs) and mTOR inhibitors—we retrained only the last layer of the finetuned VAE and added a task-specific output layer to predict OS.

In this joint training setup, each VAE was dedicated to one treatment group and trained to learn OS-related features unique to its respective cohort. Both models shared the core latent space from the unsupervised fine-tuning, allowing them to retain foundational biological representations while specializing in treatment-specific OS prediction. The training objective combined the individual OS prediction losses from each VAE into a total loss, defined as:

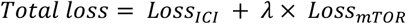

Where Loss_ICI_ and Loss_mTOR_ are the OS prediction losses for the ICI and mTOR models, respectively and λ is a weighting factor. This combined loss encouraged both models to jointly learn OS-associated features in a treatment-agnostic manner by aligning their representations in the latent space. To optimize the model, we used grid search to tune the learning rate, while the regularization parameters and the number of epochs were fixed to prevent overfitting and monitor performance metrics on validation data.

#### Adversarial Learning for Predictive Model development

To develop a treatment-specific model for predicting overall survival (OS) in response to immune checkpoint inhibitor (ICI) therapy, we implemented an Adversarial VAE model designed to isolate features unique to ICI treatment. The approach involved two VAE models trained in parallel: a primary VAE to capture OS-associated features for ICI-treated patients, and an adversarial VAE to capture OS-related features for mTOR-treated patients. Both models were initialized from the previously fine-tuned VAE, with only the last layer retrained to specialize for their respective tasks.

The main VAE was trained to predict OS for ICI patients, aiming to learn biomarkers specific to ICI response. Concurrently, the adversarial VAE was trained to predict OS for patients treated with mTOR inhibitors. The adversarial component introduced a competing objective, discouraging the main VAE from learning treatment-independent features by capturing only OS-associated patterns unique to ICI therapy. This adversarial setup was implemented by setting the total loss as the difference between the ICI-specific loss and the adversarial loss, defined as:

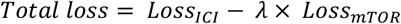

where Loss_ICI_ and Loss_mTOR_ are the OS prediction losses for the ICI and mTOR VAEs, respectively, and λ is a weighting factor that balances the adversarial effect. This formulation enables the main VAE to retain only ICI-specific features by minimizing the confounding influence of treatment-independent survival factors. Similar to joint learning, we use grid search to tune the learning rate and consider fixed regulation parameters to prevent overfitting and monitor the model performance metrics on validation data.

#### SHAP Computation

We interpreted two baseline VAEs, a prognostic VAE (outcome risk across all patients) and a predictive VAE (ICI-specific outcome risk). For each model, we computed SHAP^49^ on the held-out baseline test set using PyTorch models, shap.GradientExplainer (expected gradients) with a training-set background (fixed random subset, ∼200 samples; same background reused within each model to avoid leakage).

For a sample *x* with model output *f*(*x*), SHAP assigns feature attributions *ϕ*_*j*_(*x*)satisfying local additivity:

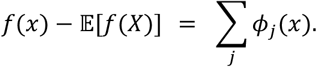

We summarized each feature *j* by importance

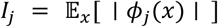

(used for ranking) and direction

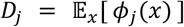

(mean signed attribution). Direction was mapped to biology according to the model’s output: if higher output means higher risk, then *D*_*j*_ < 0= protective and *D*_*j*_ > 0= harmful (reversed if the output encodes poor or favorable). We also report auxiliary summaries (SD of *ϕ*_*j*_, fraction of positive/negative attributions, and a consistency index | *D*_*j*_ |/*I*_*j*_). Top-20 lists per model were formed by sorting on *I*_*j*_. Putative metabolite annotations were for display only and did not affect SHAP computation or ranking.

#### Traditional machine-learning models

For comparison with the VAE models, we implemented traditional machine-learning approaches to perform classification and survival analysis tasks. Binary classification tasks were performed using logistic regression, with model performance evaluated using the Area Under the Curve (AUC) metric. For multi-class classification, we extended logistic regression using a one-vs-rest approach, where the model independently predicts each class against all others, creating a multi-class framework. Model performance for multi-class classification was assessed using the F1 score, which captures a balance between precision and recall.

For survival analysis, we applied the Kaplan-Meier (KM) estimator, a non-parametric method, to estimate survival functions from right-censored data. The Kaplan-Meier model provided an estimate of overall survival (OS) probabilities over time, allowing us to compare survival distributions across risk groups. Statistical significance in survival differences between groups was evaluated using the log-rank test. The concordance index and hazard ratios associated with survival predictions in different groups were evaluated by Cox proportional hazards models.

