## Supplementary information: Supplementary Tables 1-5 and Figs. 1-11 for "mzLearn, a data-driven LC/MS signal detection algorithm, enables pre-trained generative models for enhanced patient stratification"

### Supplementary Tables and Figures:

**Supplementary Table 1: Public Datasets Used for mzLearn Evaluation of Performance and Scalability**

This table summarizes the 36 publicly available datasets that evaluate mzLearn's performance and scalability. The datasets are categorized into three groups: benchmarking datasets, normalization evaluation datasets, and pretraining datasets. The table includes the number of studies in each category, the selection criteria, and data sources for each dataset. These datasets were chosen to assess signal detection quality, normalization efficiency, and the ability to develop pre-trained generative models for untargeted metabolomics.

| Public Data Sets | No. of Studies | Selection Criteria | Sources |
|---|---|---|---|
| Benchmarking datasets | 15 | Availability of targeted metabolite with reported m/z and rt values | Metabolights, Metabolomics Workbench |
| Normalization evaluation datasets | 6 | over 150 samples size with available experimental QC samples | Metabolights, Metabolomics Workbench, Mass IVE |
| Pretraining datasets | 22 | Blood-based untargeted metabolomics with the same choreography setting (i.e. HILIC positive) | Metabolomics workbench, Stanford iPOP |
| **Union** | **36** | | |

**Supplementary Table 2: Public LC–MS datasets and references.**
Study accession (Study ID), source repository, and short citation (DOI/PMID) for all datasets analyzed with mzLearn; used for benchmarking, normalization, and/or pretraining.

| Study ID | Source | Ref. article (short) |
|---|---|---|
| **MSV000086486** | Mass IVE | Delabrière et al., Anal Chem (2021). doi:10.1021/acs.analchem.1c02687. |
| **MSV000087434** | Mass IVE | Chen et al., Nat Methods (2021). doi:10.1038/s41592-021-01303-3. |
| **MTBLS1124** | Metabolights | Cai et al., Sci Rep (2020). doi:10.1038/s41598-020-61851-0. |
| **MTBLS606** | Metabolights | Shen et al., Nat Commun (2019). doi:10.1038/s41467-019-09550-x. |
| **MTBLS620** | Metabolights | Lamichhane et al., Sci Rep (2018). doi:10.1038/s41598-018-28907-8. |
| **MTBLS733** | Metabolights | Li et al., (2018). PMID:29907290. |
| **ST000387** | Metabolomics Workbench | Dev PK et al., Metabolomics (2025). |
| **ST000388** | Metabolomics workbench | Fahrmann et al., Cancer Biomark (2016). PMID:27002763. |
| **ST000422** | Metabolomics workbench | Dutta D et al., J Clin Endocrinol Metab (2016). |
| **ST000601** | Metabolomics workbench | Cruickshank-Quinn CI et al., Sci Rep (2018). |
| **ST000909** | Metabolomics workbench | Goodrich JA et al., Environ Health Perspect (2023). |
| **ST001235** | Metabolomics Workbench | Li et al., Nat Commun (2019). doi:10.1038/s41467-019-12361-9. |
| **ST001236** | Metabolomics workbench | Li et al., Nat Commun (2019). doi:10.1038/s41467-019-12361-9. |
| **ST001237** | Metabolomics Workbench | Li et al., Nat Commun (2019). doi:10.1038/s41467-019-12361-9. |
| **ST001385** | Metabolomics Workbench | No peer-reviewed article linked (CHEAR/HHEAR project record). |
| **ST001408** | Metabolomics workbench | Vykoukal et al., Nat Commun (2020). |
| **ST001422** | Metabolomics workbench | Barry et al., Cancer Prev Res (2022). doi:10.1158/1940-6207.CAPR-21-0555. |
| **ST001423** | Metabolomics workbench | Barry et al., Cancer Prev Res (2022). doi:10.1158/1940-6207.CAPR-21-0555. |
| **ST001428** | Metabolomics workbench | Jain et al., J Pediatr Gastroenterol Nutr (2024). |
| **ST001519** | Metabolomics Workbench | Tanes et al., Cell Host Microbe (2021). doi:10.1016/j.chom.2020.12.012. |
| **ST001520** | Metabolomics Workbench | McGlinchey et al., JHEP Rep (2022). doi:10.1016/j.jhepr.2022.100477. |
| **ST001710** | Metabolomics Workbench | McGlinchey et al., JHEP Rep (2022). doi:10.1016/j.jhepr.2022.100477. |
| **ST001849** | Metabolomics Workbench | Sindelar et al., Cell Rep Med (2021). doi:10.1016/j.xcrm.2021.100369. |
| **ST001918** | Metabolomics workbench | Rothman et al., Carcinogenesis (2021). doi:10.1093/carcin/bgab089. |
| **ST001931** | Metabolomics workbench | Project summary (ESPINA adolescents). |
| **ST001932** | Metabolomics workbench | Project summary (SEARCH for Diabetes in Youth). |
| **ST002027** | Metabolomics workbench | Folz JS, PhD thesis (UC Davis, 2021). |
| **ST002112** | Metabolomics workbench | Lancaster et al., Cell Host Microbe (2022). |
| **ST002233** | Metabolomics Workbench | Li S et al., Nat Commun (2023). |
| **ST002244** | Metabolomics workbench | Wang et al., medRxiv (2022). doi:10.1101/2022.09.27.22280432. |
| **ST002251** | Metabolomics workbench | Cottrill et al., J Allergy Clin Immunol (2023). doi:10.1016/j.jaci.2022.07.027. |
| **ST002331** | Metabolomics workbench | Lee et al., Hum Genomics (2022). doi:10.1186/s40246-022-00440-w. |
| **ST002711** | Metabolomics Workbench | Che et al., Mol Psychiatry (2023). doi:10.1038/s41380-023-02051-w. |
| **ST002773** | Metabolomics workbench | Rahman et al., Int J Cancer (2024). doi:10.1002/ijc.34929. |
| **ST002817** | Metabolomics Workbench | Vaniya et al., bioRxiv (2023). doi:10.1101/2023.11.10.564562. |
| **Stanford-hmp2** | http://hmp2-data.stanford.edu/ | iHMP Consortium, Nature (2019). doi:10.1038/s41586-019-1238-8. |

**Supplementary Table 3: mzLearn outperforms existing LC/MS signal detection methods.** We evaluated mzLearn's performance against existing signal detection methods using 15 publicly available LC/MS datasets acquired with diverse LC/MS instrumentation. mzLearn consistently demonstrated a higher true-positive rate and a lower false-positive rate compared to ASARI and XCMS. Additionally, mzLearn captured, on average, the highest number of detected signals with a minimum of 20% frequency.

| | | | | | **Average of number of peaks > 20% frequency** | | | **True Positive Rate** | | | **False Positive Rate** | | |
|---|---|---|---|---|---|---|---|---|---|---|---|---|---|
| **Study ID** | **Source** | **LC type** | **Ion** | **MS type** | **ASARI** | **XCMS** | **mzLearn** | **ASAR** | **XCMS** | **mzLearn** | **ASARI** | **XCMS** | **mzLearn** |
| ST002711 | Metabolomics Workbench | HILIC | pos | QTOF | 167 | 1,225 | 4,802 | 20% | 57% | 84% | 25% | 15% | 10% |
| ST001519 | Metabolomics Workbench | RP | neg | Orbitrap | 10,591 | 14,622 | 14,423 | 78% | 89% | 93% | 15% | 23% | 10% |
| ST001519 | Metabolomics Workbench | HILIC | neg | Orbitrap | 4,829 | 4,442 | 4,871 | 71% | 82% | 85% | 37% | 16% | 18% |
| ST000387 | Metabolomics Workbench | RP | pos | QTOF | 245 | 1,392 | 8,788 | 23% | 80% | 95% | 8% | 4% | 22% |
| ST002233 | Metabolomics Workbench | HILIC | pos | Orbitrap | 8,288 | 3,197 | 8,149 | 85% | 33% | 82% | 22% | 48% | 7% |
| ST001710 | Metabolomics Workbench | RP | pos | QTOF | - | 6,570 | 10,118 | - | 81% | 80% | - | 11% | 8% |
| MTBLS620 | Metabolights | RP | pos | QTOF | 581 | 4,688 | 12,018 | 39% | 81% | 90% | 3% | 9% | 9% |
| ST001235 | Metabolomics Workbench | HILIC | pos | Triple quadrupole | 9,204 | 7,980 | 10,720 | 79% | 91% | 92% | 23% | 12% | 13% |
| ST001237 | Metabolomics Workbench | HILIC | pos | Orbitrap | 7,520 | 7,648 | 10,272 | 79% | 95% | 98% | 20% | 22% | 9% |
| MTBLS606 | Metabolights | HILIC | neg | QTOF | 269 | 3,708 | 8,784 | 0% | 24% | 74% | 28% | 29% | 26% |
| ST001385 | Metabolomics Workbench | HILIC | pos | QTOF | 473 | 7,659 | 11,900 | 31% | 79% | 89% | 19% | 8% | 10% |
| ST002817 | Metabolomics Workbench | RP | neg | QTOF | 213 | 2,340 | 9,146 | 23% | 81% | 92% | 8% | 17% | 13% |
| ST001849 | Metabolomics Workbench | HILIC | pos | QTOF | 500 | 2,816 | 6,266 | 46% | 80% | 89% | 17% | 11% | 15% |
| MTBLS1124 | Metabolights | HILIC | neg | QTOF | 1,028 | 9,753 | 21,032 | 13% | 90% | 87% | 23% | 29% | 17% |
| MSV000087434 | Mass IVE | HILIC | neg | Orbitrap | 3,704 | 8,086 | 6,174 | 94% | 98% | 96% | 6% | 16% | 11% |
| MTBLS733 | Metabolights | RP | pos | Orbitrap | 22,209 | 27,478 | 35,615 | 42% | 97% | 98% | 28% | 6% | 2% |
| **Average** | | | | | 4,654.73 | 7,100.25 | **11,442.38** | 48.2% | 77.4% | **89.0%** | 18.8% | 17.3% | **12.5%** |

**Supplement Table 4: mzLearn overcomes signal intensity drifts even in the absence of QC samples.**

We benchmarked mzLearn's ability to address signal intensity drifts caused by run order and batch effects with and without QC samples. Six publicly available, large-scale datasets with QC samples were used for validation. We concealed 10% of pooled QC samples, applied various normalization methods, and calculated the relative standard deviation (RSD) of the hidden pooled samples. The table below compares RSD for raw data, total ion intensity normalization, QC-based normalization, and synthetic QC-based normalization, with the synthetic QC method performing comparably to QC-based normalization in mitigating signal drifts.

| | | | | | | | | **RSD of hidden pool samples** | | | |
|---|---|---|---|---|---|---|---|---|---|---|---|
| **Study ID** | **Source Database** | **LC type** | **Ion** | **No. Peaks** | **No. Samples** | **No. Pool Samples** | **No. batches** | **Raw** | **TIC Normalization** | **QC-based Normalization** | **Synthetic QC-based Normalization** |
| MTBLS1124 | Metabolights | HILIC | Negative | 11,702 | 264 | 28 | 1 | 32% | 23% | **18%** | 19% |
| MSV000086486 | Mass IVE | RP | Positive | 2,397 | 2,386 | 81 | 2 | 20% | 20% | **13%** | 16% |
| ST000387 | Metabolomics Workbecnh | RP | Positive | 8,570 | 276 | 26 | 1 | 14% | 13% | **12%** | 12% |
| ST002233 | Metabolomics Workbecnh | HILIC | Positive | 4,810 | 184 | 19 | 14 | 44% | 44% | 34% | **33%** |
| ST001520 | Metabolomics Workbecnh | RP | Negative | 2,862 | 177 | 25 | 1 | 19% | 19% | **10%** | 11% |
| ST001237 | Metabolomics Workbecnh | HILIC | Positive | 4,537 | 1,379 | 132 | 2 | 22% | 18% | **12%** | 13% |
| **Average** | | | | | | | | **25%** | **23%** | **16%** | **17%** |

**Supplement Table 5: Pretrain datasets.**
The 22 datasets used for pre-training, along with their associated IDs, the type of LC column, ion mode, the number of samples per study, and the number of peaks detected by mzLearn in at least 20% of the study samples.

| Study ID | Source | LC type | Ion | Cohort No. | Number of Samples | Number of Peaks |
|---|---|---|---|---|---|---|
| ST002773 | Metabolomics workbench | HILIC | Positive | 1 | 2963 | 2,541 |
| ST002331 | Metabolomics workbench | HILIC | positive | 2 | 1315 | 6,079 |
| ST002251 | Metabolomics workbench | HILIC | positive | 3 | 210 | 7,992 |
| ST002244 | Metabolomics workbench | HILIC | positive | 4 | 122 | 11,816 |
| ST002112 | Metabolomics workbench | HILIC | positive | 5 | 335 | 14,677 |
| ST002027 | Metabolomics workbench | HILIC | positive | 6 | 356 | 3,187 |
| ST001932 | Metabolomics workbench | HILIC | positive | 7 | 4482 | 46,954 |
| ST001931 | Metabolomics workbench | HILIC | positive | 8 | 2044 | 5,354 |
| ST001918 | Metabolomics workbench | HILIC | positive | 9 | 217 | 8,979 |
| ST001849 | Metabolomics workbench | HILIC | positive | 10 | 691 | 6,233 |
| ST001519 | Metabolomics workbench | HILIC | positive | 11 | 148 | 9,656 |
| ST001428 | Metabolomics workbench | HILIC | positive | 12 | 1522 | 8,207 |
| ST001423 | Metabolomics workbench | HILIC | positive | 13 | 1080 | 8,422 |
| ST001422 | Metabolomics workbench | HILIC | positive | 14 | 1799 | 10,861 |
| ST001408 | Metabolomics workbench | HILIC | positive | 15 | 349 | 11,793 |
| ST001237 | Metabolomics workbench | HILIC | positive | 16 | 638 | 9,778 |
| ST001236 | Metabolomics workbench | HILIC | positive | 17 | 271 | 9,778 |
| ST000909 | Metabolomics workbench | HILIC | positive | 18 | 742 | 6,865 |
| ST000601 | Metabolomics workbench | HILIC | positive | 19 | 384 | 3,596 |
| ST000422 | Metabolomics workbench | HILIC | positive | 20 | 60 | 4,214 |
| ST000388 | Metabolomics workbench | HILIC | positive | 21 | 94 | 26,549 |
| Stanford-iPOP | http://hmp2-data.stanford.edu/ | HILIC | positive | 22 | 726 | 17,143 |
| **Sum** | | | | | **20,548** | **240,674** |

**Supplement Table 6: Fine-tune datasets and meta-data splits**

Clinical data for the CheckMate 025 Phase III clinical trial of clear cell-renal cell carcinoma patients undergoing immune-checkpoint inhibitor (ICI) therapy (nivolumab, n=392) or mTOR inhibition (everolimus, n=349). Samples were split into 60% training, 20% validation, and 20% test sets. We calculated chi-square (for categorical variables) and ANOVA (for continuous survival variables) statistics and p-values to ensure the distribution of clinical variables was not different between training, validation, and test sets.

| | | NIVOLUMAB | | | | EVEROLIMUS | | | |
|---|---|---|---|---|---|---|---|---|---|
| | | **training (n=234)** | **validation (n=79)** | **test (n=79)** | **p** | **training (n=209)** | **validation (n=70)** | **test (n=70)** | **p** |
| **Benefit** | CB | 71 (30.3%) | 24 (30.4%) | 26 (32.9%) | 0.994 | 56 (26.8%) | 19 (27.1%) | 20 (28.6%) | 0.998 |
| | ICB | 84 (35.9%) | 28 (35.4%) | 28 (35.4%) | | 99 (47.4%) | 33 (47.1%) | 33 (47.1%) | |
| | NCB | 79 (33.8%) | 27 (34.2%) | 25 (31.6%) | | 54 (25.8%) | 18 (25.7%) | 17 (24.3%) | |
| **Sex** | Female | 56 (23.9%) | 19 (24.1%) | 19 (24.1%) | 1 | 56 (26.8%) | 19 (27.1%) | 22 (31.4%) | 0.748 |
| | Male | 178 (76.1%) | 60 (75.9%) | 60 (75.9%) | | 153 (73.2%) | 51 (72.9%) | 48 (68.6%) | |
| **Age group** | >75 | 19 (8.1%) | 7 (8.9%) | 6 (7.6%) | 0.975 | 21 (10.0%) | 9 (12.9%) | 6 (8.6%) | 0.893 |
| | 65-75 | 66 (28.2%) | 24 (30.4%) | 25 (31.6%) | | 68 (32.5%) | 20 (28.6%) | 21 (30.0%) | |
| | <65 | 149 (63.7%) | 48 (60.8%) | 48 (60.8%) | | 120 (57.4%) | 41 (58.6%) | 43 (61.4%) | |
| **Region** | REST OF WORLD | 56 (23.9%) | 15 (19.0%) | 20 (25.3%) | 0.874 | 50 (23.9%) | 18 (25.7%) | 20 (28.6%) | 0.865 |
| | US/CANADA | 99 (42.3%) | 37 (46.8%) | 32 (40.5%) | | 89 (42.6%) | 26 (37.1%) | 26 (37.1%) | |
| | WESTERN EUROPE | 79 (33.8%) | 27 (34.2%) | 27 (34.2%) | | 70 (33.5%) | 26 (37.1%) | 24 (34.3%) | |
| **MSKCC** | FAVORABLE | 80 (34.2%) | 27 (34.2%) | 27 (34.2%) | 1 | 70 (33.5%) | 27 (38.6%) | 26 (37.1%) | 0.703 |
| | INTERMEDIATE | 109 (46.6%) | 37 (46.8%) | 36 (45.6%) | | 104 (49.8%) | 30 (42.9%) | 29 (41.4%) | |
| | POOR | 45 (19.2%) | 15 (19.0%) | 16 (20.3%) | | 35 (16.7%) | 13 (18.6%) | 15 (21.4%) | |
| **IMDC** | FAVORABLE | 30 (13.3%) | 12 (15.8%) | 13 (17.1%) | 0.804 | 39 (19.5%) | 10 (14.9%) | 10 (14.9%) | 0.29 |
| | INTERMEDIATE | 143 (63.3%) | 43 (56.6%) | 46 (60.5%) | | 128 (64.0%) | 39 (58.2%) | 40 (59.7%) | |
| | POOR | 53 (23.5%) | 21 (27.6%) | 17 (22.4%) | | 23 (16.5%) | 18 (26.9%) | 17 (25.4%) | |
| **Prior antiangiogenic treatments >1** | FALSE | 181 (77.4%) | 61 (77.2%) | 61 (77.2%) | 1 | 158 (75.6%) | 55 (78.6%) | 54 (77.1%) | 0.87 |
| | TRUE | 53 (22.6%) | 18 (22.8%) | 18 (22.8%) | | 51 (24.4%) | 15 (21.4%) | 16 (22.9%) | |
| **OS event** | No | 70 (29.9%) | 23 (29.1%) | 25 (31.6%) | 0.937 | 44 (21.1%) | 14 (20.0%) | 14 (20.0%) | 0.972 |
| | Yes | 164 (70.1%) | 56 (70.9%) | 54 (68.4%) | | 165 (78.9%) | 56 (80.0%) | 56 (80.0%) | |
| **PFS event** | No | 32 (13.7%) | 9 (11.4%) | 10 (12.7%) | 0.868 | 28 (13.4%) | 9 (12.9%) | 9 (12.9%) | 0.989 |
| | Yes | 202 (86.3%) | 70 (88.6%) | 69 (87.3%) | | 181 (86.6%) | 61 (87.1%) | 61 (87.1%) | |
| **OS** | mean (std) | 29.7 (19.8) | 28.9 (20.9) | 30.8 (20.6) | 0.835 | 25.6 (19.9) | 25.3 (18.6) | 25.4 (19.6) | 0.994 |
| **PFS** | mean (std) | 9.2 (13.0) | 9.6 (13.7) | 9.0 (13.3) | 0.957 | 6.8 (8.1) | 6.3 (6.6) | 7.3 (9.2) | 0.767 |

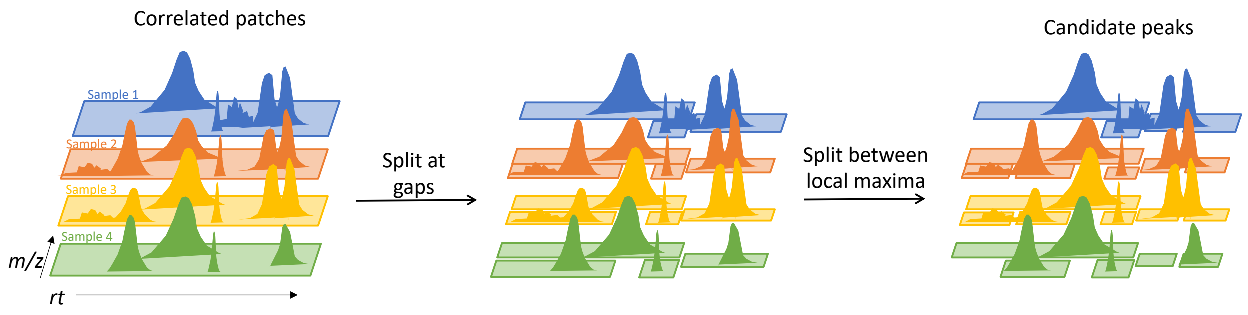


**Supplementary Figure 1: Scissor Operation.** After the pairwise file cross-correlation procedure, overlapping correlated patch groups are merged, which may result in multiple peaks sharing a single patch-group. The figure shows an example of correlated patches across four samples containing multiple signals (left). The first step in the scissor operation splits the patch at the gaps in *rt* and *m/z* space, which in the above example creates four correlated patch-groups (center). The next step in the scissor operation splits patches at the valleys between local intensity maxima (right). In the above example, this results in a total of six correlated patch-groups. The resulting patch-groups are now candidate peaks, which become proper peak-groups (features) after filtering out the low-quality candidate peaks.

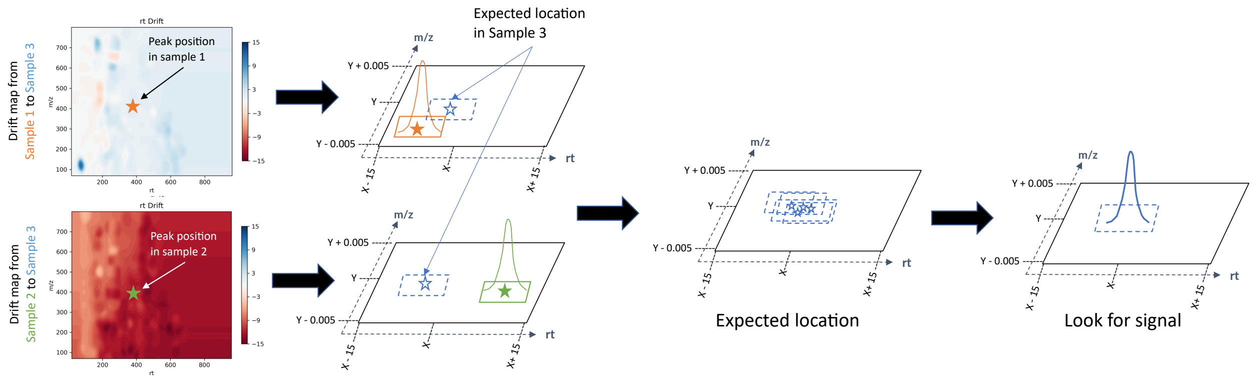


**Supplementary Figure 2: Missing Peak Search.** The missing peak search rescues signals previously missed or not captured due to the application of earlier quality filters or less accurate drift maps in prior iterations. The figure illustrates an example where the drift map from sample 1 to sample 3 provides an initial estimate for the expected peak location in sample 3. Similarly, the drift map from sample 2 to sample 3 offers a second estimate for the same peak. By combining these estimates, we calculate an average expected location for the peak in sample 3. This location is then used to search for potential missing signals. If a signal is identified in sample 3 and meets the updated quality criteria, it is incorporated into the existing peak groups

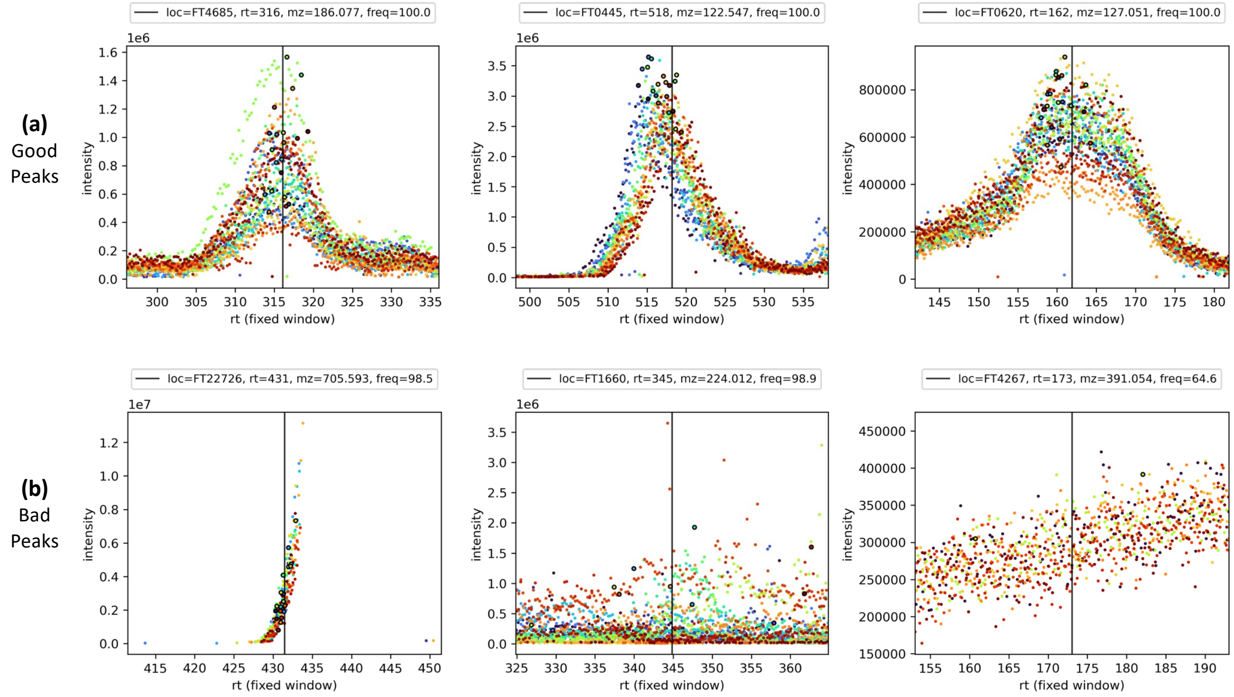


**Supplementary Figure 3: Examples of Graded Peaks from the ST001236 data set. (a)** "Good" peaks were those signals with a clear bell shape, correctly reported signal maximum intensity, and aligned signals across samples. **(b) "**Bad" peaks had questionable or irregular shape and/or misalignment issues as judged by the recorded locations of maximum intensity. Peaks from 20 samples are plotted, with the different colors referring to different samples. To improve visibility, only 10 colors are used, and each color refers to a signal from up to two different sample files. The vertical black line indicates the identified median rt of the peak. The data points indicate the coordinates for a sample's recorded maximum intensity with the black outline.

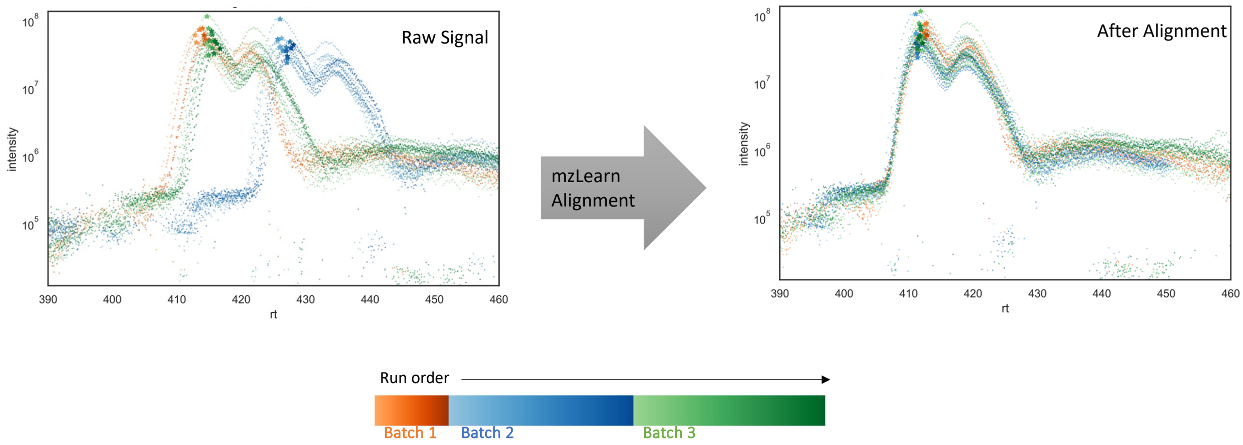


**Supplementary Figure 4: mzLearn can correctly estimate the rt drift between leucine and iso-leucine peaks.**
Peaks from the three batches in the combined ST001236 and ST001237 datasets initially show rt drift due to run order and batch effects. After mzLearn alignment, the peaks are correctly aligned, with iso-leucine peak locations highlighted at the start. For clarity, 2–3 randomly selected samples are plotted from every 50 files.

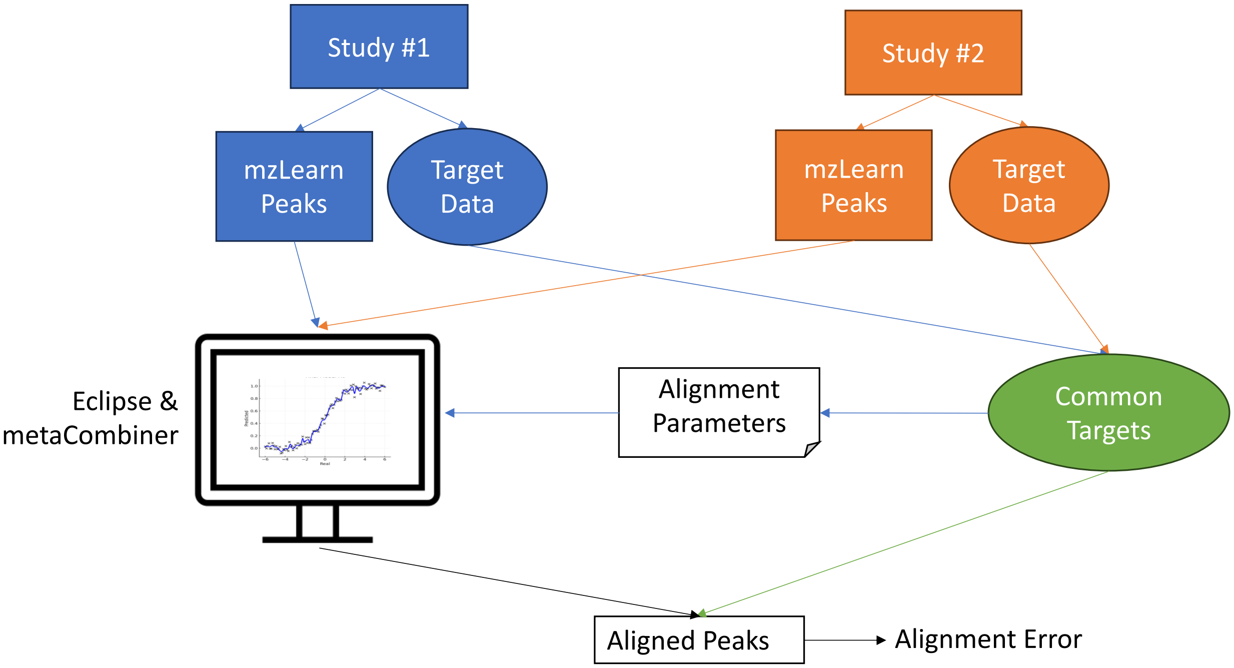


**Supplementary Figure 5: aligning the peaks between two independent studies.**
To align peaks between two independent studies, we first identify targeted peaks by name matching. The differences in m/z and rt values for these targeted peaks are then used to inform alignment parameters. Two retention time (rt) alignment methods, Eclipse and metaCombiner, are applied to project the rt values of both targeted and untargeted features. The union of these results is used as the final set of aligned peaks. Comparing these aligned results with the targeted data alignment allows us to calculate the alignment error and assess accuracy.

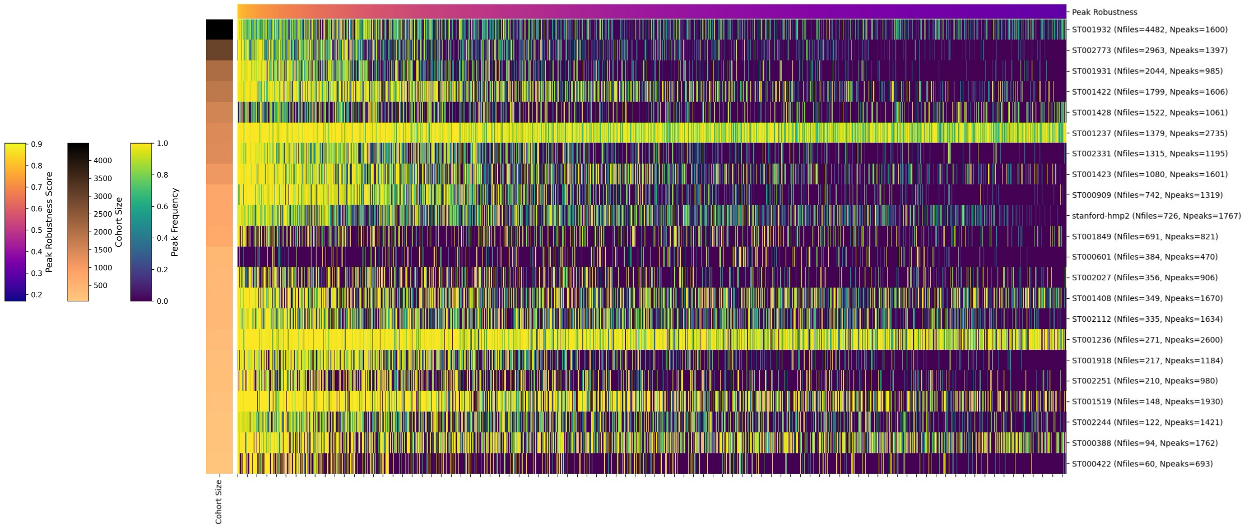


**Supplementary Figure 6: A heat map of peak frequencies and robustness scores across pretraining and fine-tuning datasets.**

Following peak alignment across pretraining and fine-tuning datasets, we calculated a robustness score for each peak to assess its detection consistency across studies. The figure presents a heatmap displaying peak frequencies across studies. The heatmap illustrates peak frequencies across studies, with yellow indicating a frequency of 1 and dark blue indicating a frequency of 0. The top bar graph displays the corresponding robustness scores for each peak.

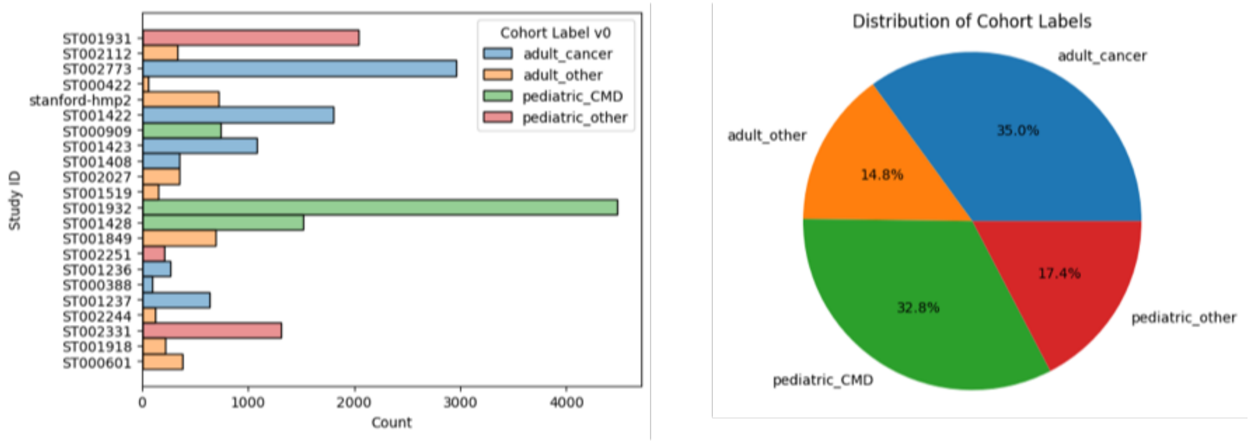


**Supplementary Figure 7: Pretraining datasets and associated labels.**
While exact metadata was unavailable for all pretraining samples, we categorized them based on available information. Samples were labeled as adult or pediatric studies and further classified as cancer, cardio-metabolic (CMD), or other when relevant.

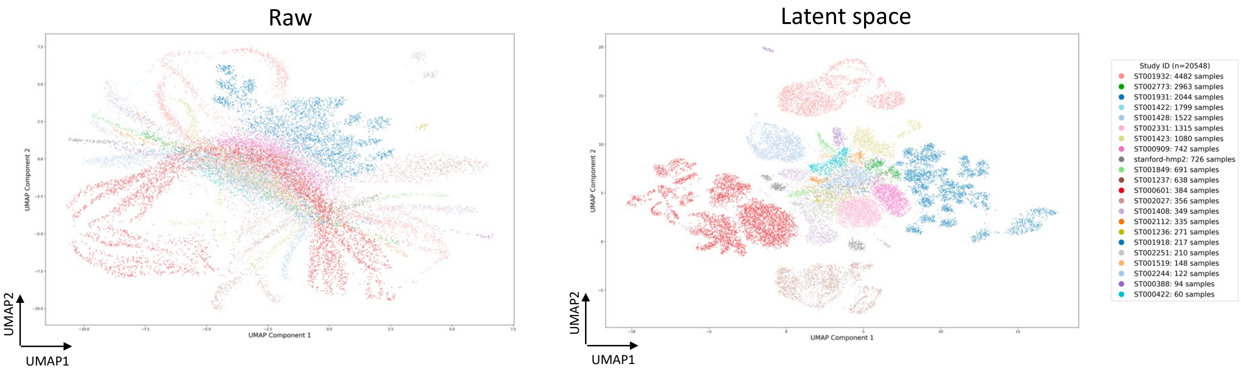


**Supplementary Figure 8: Raw Low-Dimensional Representation of pre-trained VAE Models based color-coded by study ID.**

The UMAP visualization of raw datasets shows the mixing of the samples by study IDs. As the studies encompass subjects with varying disease conditions and age groups, the latent space of the pre-trained VAE models further separates samples based on their underlying disease and age characteristics.

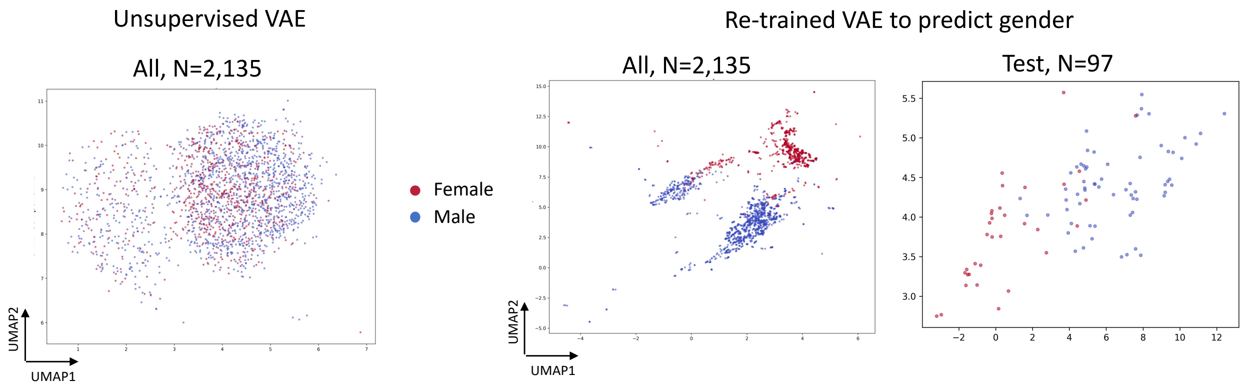


**Supplementary Figure 8: Low-Dimensional Representation of Pre-Trained VAE Models Before and After Retraining for Gender Prediction**
The low-dimensional representation of an unsupervised VAE model is depicted before (left panel) and after (right panel) supervised retraining for gender prediction. Initially, the models were developed in an unsupervised manner, capturing latent patterns in the data that do not distinctly separate genders. During retraining, only the final layers of the encoder from the pre-trained models were fine-tuned using gender labels. This retraining significantly improved the model's ability to differentiate gender, achieving an AUC of 96.02% on the unseen test set.

### Transfer Learning

#### Best Trial

Trial number: 38

Objective value: 0.8418963074684143

Best Parameters:
dropout_rate: 0.15000000000000002
learning_rate: 0.000136159422872435
l1_reg: 2.3618027973370643e-06
weight_decay: 0.0008931324480984989
num_epochs: 40

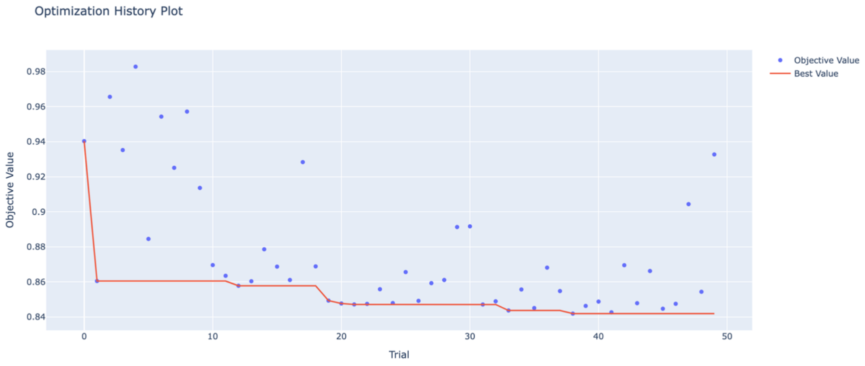


### Random initialized

#### Best Trial

Trial number: 43

Objective value: 0.9629389762878418

Best Parameters:
dropout_rate: 0.1
learning_rate: 0.00047788791057886033
l1_reg: 1.4267287743774265e-06
weight_decay: 1.0285521899983445e-06
num_epochs: 50

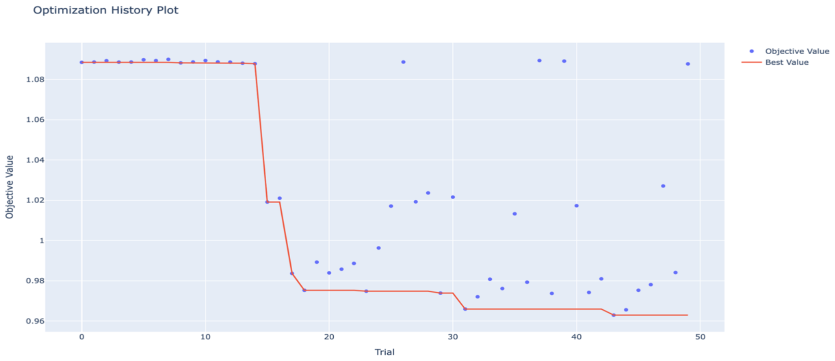


**Supplementary Figure 10: Optuna Trials for Fine-Tuned VAE Models with and without Transfer Learning.**
Optuna trials comparing fine-tuned VAE models with transfer learning to randomly initialized models. Models with transfer learning required fewer trials to achieve optimal performance and reached a lower validation loss of 0.84 compared to 0.96 for randomly initialized models.

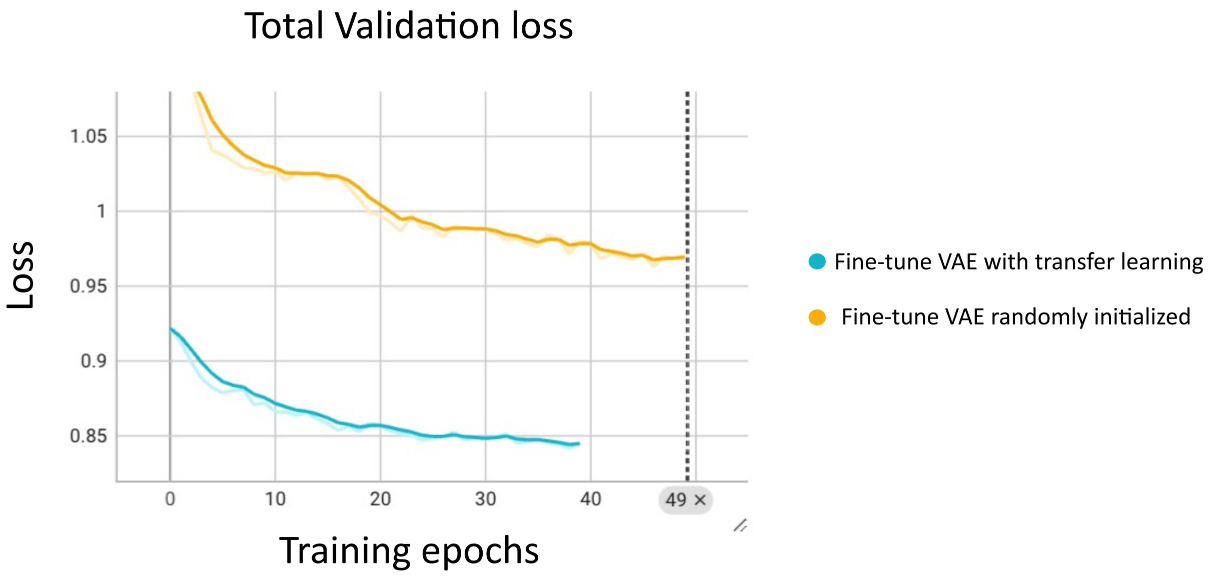


**Supplementary Figure 11: Validation Loss Across Training Epochs for Fine-Tuned VAE Models.**

Validation loss during training epochs for fine-tuned VAE models with transfer learning compared to randomly initialized models. For the selected optimal parameters using Optuna, VAE models with transfer learning start with a lower initial loss, reach the minimum loss in fewer epochs, and achieve a lower final loss than randomly initialized models.
